# Human lipidomics guides brain-inspired lipoprotein particle approach to ameliorate Alzheimer’s disease pathology

**DOI:** 10.64898/2026.11.25.769060

**Authors:** Rebecca L. Pinals, Yeojung Koh, Ziying Ke, Anjanet Tuyéras, Claudia F. Lozano Cruz, Ilyas Gajarski, Aitana Burman, Cheng-Yi Yang, Arthur Ngo, Lucas B. Choy, Jake Weiss, Pavitra Balachundhar, Daniel Leible, Robert Hurt, Aaron Choi, Ivette Chen, Eulim Kang, Djuna von Maydell, Leyla A. Akay, Daniela Rodrigues-Amorim, Erica McNamara, Kyriaki Galani, James Cameron, Florence Liang, Amy Deik, Kerry Pierce, DongSoo Yun, David A. Bennett, Clary Clish, Li-Huei Tsai

## Abstract

*APOE* encodes the principal lipid-transporting protein in the brain, and lipid dysregulation is increasingly implicated in Alzheimer’s disease (AD) pathogenesis.^1,2^ The *APOE4* allele, together with advanced age and female sex, are major risk factors for late-onset AD.^3^ Yet how these risk factors interact to reshape lipid composition and transport in the brain remains unclear. Herein, we characterize lipidomic profiles in the brain, cerebrospinal fluid (CSF), and blood of individuals with or without AD, as well as of mice expressing humanized *APOE*. We identify brain lipidomic shifts associated with AD in humans, including relative depletion of highly unsaturated lipids, with many changes resembling those associated with *APOE4* alone. Individuals with AD show striking depletion of lipoprotein particle (LP)-associated lipid abundance in the CSF. Additionally, humanized *APOE4* mice exhibit an age-dependent deficit in CSF LP-associated lipid abundance relative to *APOE3* mice. Accordingly, we leverage these insights to engineer brain-inspired reconstituted LPs (rLPs) with the goal of restoring lipid transport in the brain. rLPs promote cholesterol efflux from human induced pluripotent stem cell-derived *APOE4* astrocytes *in vitro*. In an AD mouse model (5xFAD) expressing humanized *APOE4*, these nanoparticles reduce lipid-droplet accumulation, facilitate amyloid-β clearance, enhance synaptic integrity, and improve cognitive function. Together, our results provide a resource mapping lipidomic changes across the Alzheimer’s risk triad and demonstrate that engineered rLPs can counter lipid accumulation and associated pathology in the brain, and cognitive deficits *in vivo*.

## Introduction

Amyloid-β plaques and neurofibrillary tau tangles remain hallmarks of Alzheimer’s disease (AD) pathology, yet lipid dysregulation and accumulation are also longstanding features of the disease.^4,5^ Despite this, lipids remain relatively understudied as a biomolecular class implicated in AD.^6^ Genome-wide association studies (GWAS) have underscored the role of lipid biology in AD, with many of the strongest risk loci for sporadic AD converging on lipid handling and metabolism.^7,8^ Most notably among them is *APOE*, which encodes the lipid-transporting protein, apolipoprotein E (ApoE).^1,9^ In the brain, ApoE is the predominant protein component of lipoprotein particles (LPs) that shuttle lipids between glia and neurons to support membrane synthesis and remodeling, synaptic function, and overall lipid redistribution and homeostasis.^1,10^ Three common *APOE* alleles shape AD risk in distinct ways: *APOE3* is the most prevalent and considered risk-neutral, *APOE2* is less common but protective, and *APOE4* is the strongest genetic risk factor for late-onset AD.^11–13^ Although differing from ApoE3 by only a single amino acid, the ApoE4 variant has an altered protein structure that affects its lipid-binding capacity and overall lipid transport function.^9,14,15^ Prior studies have demonstrated that *APOE4* expression disrupts lipid homeostasis, leads to the accumulation of lipid droplets in glia, and undermines essential glial support of neurons.^16,9,17–21^

The *APOE4* allele, together with advanced age and female sex, comprises the Alzheimer’s risk triad.^3^ Age remains the strongest risk factor for AD, with prevalence increasing from 5.2% among individuals aged 65–74 up to 35.8% among those aged 85 and older.^22^ Additionally, nearly two-thirds of AD patients are women,^22^ a disparity that cannot be fully explained by their greater longevity compared to men.^23^ The intertwined effects of age, *APOE*, and sex on lipid homeostasis remain poorly understood.

To understand how AD impacts lipid composition and distribution, we mapped lipid profiles in the brain, cerebrospinal fluid (CSF), and blood of both human and mouse cohorts via mass spectrometry-based lipidomics. Our results reveal that AD and its major risk factors – aging, *APOE4*, and female sex – contribute to broad disruptions in brain lipid homeostasis and transport capacity. Based on these findings, we designed, synthesized, and validated a brain-inspired formulation of reconstituted lipoprotein particles (rLPs) to counter disrupted lipid transport in AD-relevant models. Specifically, rLP treatment promoted cholesterol efflux from human induced pluripotent stem cell (iPSC)-derived *APOE4* astrocytes. In *APOE4*;5xFAD mice, rLP treatment reduced lipid-droplet accumulation and amyloid-β pathology, enhanced synaptic organization and neurite-development programs, and improved recognition memory. These findings establish rLPs as a tool to modulate brain biomolecular transport and a potential therapeutic platform for mitigating Alzheimer’s pathologies.

## Results

### Lipidomic profiling of the human MAP cohort

To investigate the effects of AD on lipidomic profiles, we analyzed two sample sets from the Rush Memory and Aging Project (MAP) for a total of 68 individuals with donor-matched postmortem prefrontal cortex (PFC) and ventricular cerebrospinal fluid (CSF) (**Figure 1a**).^24^ The primary set (*n* = 40) was balanced across AD status (pathologic diagnosis by NIA-Reagan criteria), *APOE3/3* vs. *APOE3/4* genotype, and female vs. male sex, and additionally included blood serum collected at the last visit. The second set (*n* = 28) included the same comparison groups except comprising all females. Donors were 72–100 years old at the time of death and postmortem intervals were ≤ 10 h (**Extended Data Figure 1a–b**). AD and non-AD control individuals were selected based on National Institute on Aging (NIA)-Reagan Diagnosis criteria of AD (**Extended Data Figure 1c**). Additional cohort characteristics are provided (**Extended Data Figure 1d–g**). We performed mass spectrometry-based lipidomics on each sample and identified 837 unique lipids representing 25 lipid classes across all samples after quality-control filtering. Bridge samples included across batches showed concordance (**Extended Data Figure 1h–j**), and samples were combined into a single cohort. Full details on the cohort design, tissue and biofluid preparations, and lipidomic analyses are included in Methods.

**Figure 1.**
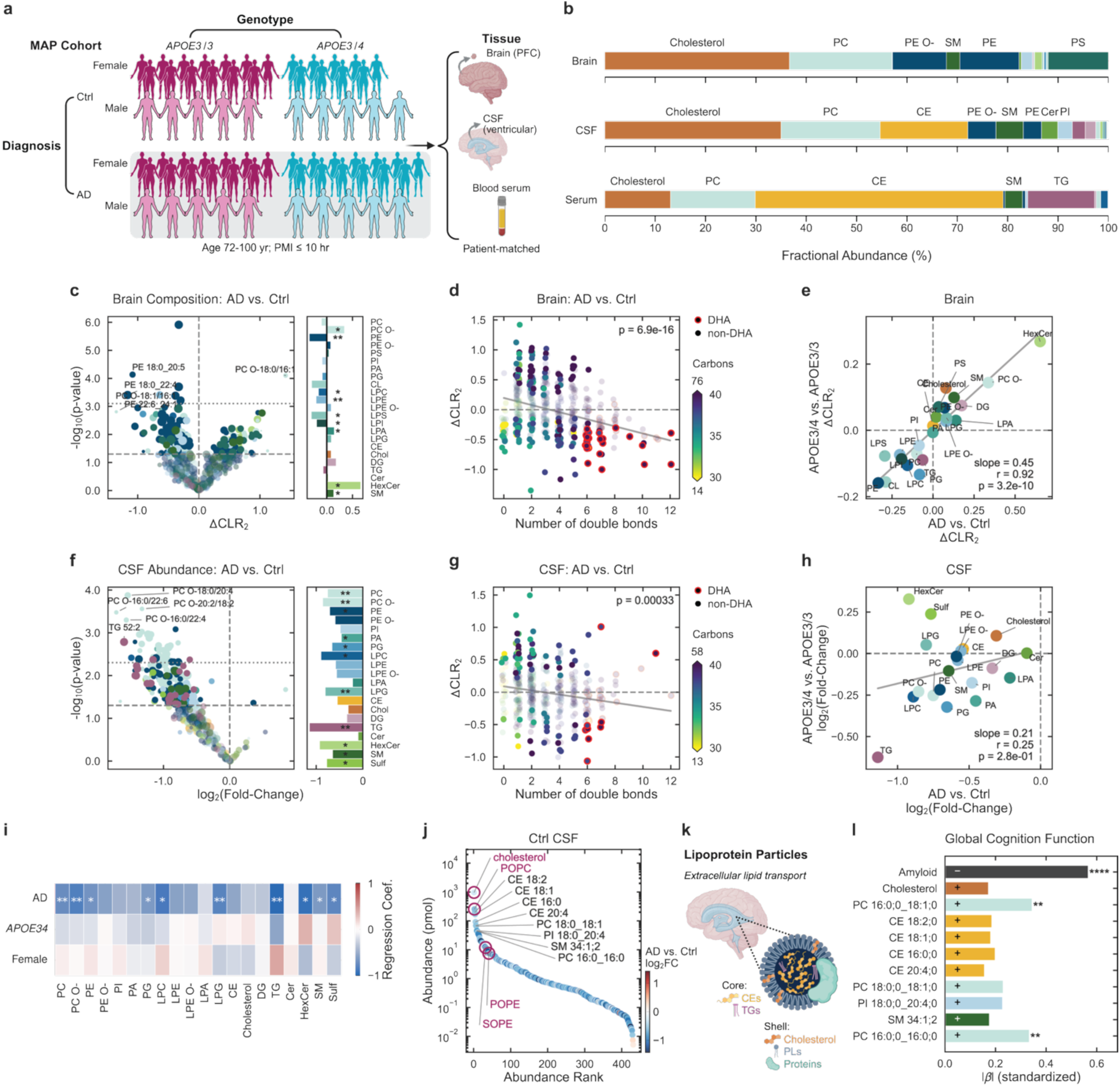
Alzheimer’s disease is associated with brain lipid compositional changes and CSF lipid depletion. **(a)** Overview of human study design with the Rush Memory and Aging Project (MAP) to compare lipid changes in individuals with (AD) or without (Ctrl) neuropathologic AD, *APOE3/3* (magenta) and *APOE3/4* (cyan) genotypes, for female (dark) and male (light) individuals (*n* = 68 total). Postmortem prefrontal cortex (PFC), ventricular cerebrospinal fluid (CSF), and blood serum collected at last visit matched per individual were analyzed by mass spectrometry-based lipidomics. Brain samples were analyzed for 32 AD and 32 Ctrl, and CSF samples were analyzed for 36 AD and 31 Ctrl individuals. **(b)** Fractional contribution of each lipid class to total measured lipid content in human brain (top), CSF (middle), and serum (bottom; only from primary set), calculated as the median of the sum of lipid abundances within each class across individuals, normalized to the total, and ordered by class abundance in CSF. **(c)** Volcano plot (-log_10_ *P* value vs. centered log_2_ ratio (CLR_2_) difference) of brain lipids in AD vs. Ctrl. Right panel bar plot shows corresponding class-level differences in CLR_2_. Markers indicate individual lipid species colored by lipid class, with size scaled by abundance and semi-transparency for *P* > 0.05. Short-dashed line indicates FDR-adjusted *P* = 0.05 (Benjamini-Hochberg correction). **(d)** Difference in CLR_2_ lipid abundance for AD vs. Ctrl brain as a function of total number of double bonds. Markers indicate individual lipid species and are colored by total carbon chain number; DHA-containing lipid markers are outlined red. Significantly changing lipids are shown with darker opacity. Gray line denotes the linear fit across all lipids, with annotated *P* value. **(e)** Class-level CLR_2_ for *APOE3/4* vs. *APOE3/3* against that of AD vs. Ctrl in the brain. Gray line denotes the linear fit across all lipid classes, with annotated slope, Pearson’s *r*, and *P* value. **(f)** As described in **(c)**, but for CSF lipids in AD vs. Ctrl and showing log_2_ fold-change in absolute lipid abundance rather than CLR_2_ difference. **(g)** As described in **(d)**, but for CSF lipids in AD vs. Ctrl. **(h)** as described in **(e)**, but for CSF lipids on log_2_ fold-change scale. **(i)** Heatmap of coefficients from multivariable regressions of log_2_-transformed summed CSF lipid abundance across a lipid class per individual. Each column represents a regression per lipid class using a linear model including terms for AD, *APOE3/4*, and female sex. *P* values for regression coefficients were determined using two-sided t-tests of coefficients from ordinary least-squares regression. **(j)** Rank-abundance plot of lipids in Ctrl CSF, with marker color denoting log_2_ fold-change in abundance for AD vs. Ctrl. Lipids are ranked by median abundance in Ctrl CSF; the ten most abundant lipids are labeled. Lipids selected for downstream studies are highlighted with magenta circles and labels. **(k)** Schematic of CSF lipoprotein particles (LPs), with lipids in the shell (phospholipids and cholesterol) and core (predominantly CEs, with some TGs), and ApoE as the primary apolipoprotein. **(l)** Absolute value of regression coefficients of global cognition against amyloid (top row; for 52 individuals for which amyloid was reported) and top 10 abundant CSF lipid species, all z-score normalized. Negative (-) and positive (+) regression coefficients are denoted on each row. Amyloid content in the brain represents the percent area of cortex occupied by amyloid measured by immunohistochemistry and averaged across eight brain regions.^30^ For statistical comparisons, linear models were used to control for sample set and nominal *P* values were determined using a two-sided t-test on the regression coefficients. * *P* < 0.05, ** *P* < 0.01, *** *P* < 0.001, **** *P* < 0.0001.

Integrated analysis of lipid species shared across the brain, CSF, and serum revealed strong compartment-specific organization: Lipid abundance profiles were most strongly correlated among samples within a given tissue or biofluid (**Extended Data Figure 2a**). Cross-sample similarity was intermediate for brain-CSF and CSF-serum, whereas brain-serum correlations were lowest, suggesting CSF may provide an intermediate biochemical interface between brain and peripheral lipid pools. From principal component analysis (PCA), CSF lipids exhibited more spread as compared to the tightly clustered brain and serum samples (**Extended Data Figure 2b**). These separations illustrate compartmentalization of lipid composition and metabolism across these biofluids and brain tissue.

Next, we examined all lipids identified within each tissue (**Figure 1b**). From the fractional contribution of each lipid class to the total lipid content, the brain lipidome was dominated by cholesterol, followed by abundant phospholipid classes including phosphatidylethanolamine (PE), ether-linked PE (PE O-), phosphatidylcholine (PC), and phosphatidylserine (PS), and the brain-abundant sphingomyelin (SM). The CSF contained similar lipid classes, though with a higher relative proportion of cholesteryl ester (CE) and ceramide (Cer) compared to the brain. In serum, CE accounted for nearly half of all lipids, while triacylglycerol (TG) was much more abundant than in brain and CSF, reflecting the distinct lipoprotein particle profiles of the central nervous system and periphery.^1^ The full landscapes of lipids as a function of AD, *APOE* genotype, and chromosomal sex are included for the brain (**Extended Data Figure 2c**), CSF (**Extended Data Figure 2d**), and serum (**Extended Data Figure 2e**).

### AD and *APOE4* modulate brain and CSF lipidome

We first analyzed lipidomic profiles in the human brain using a centered log_2_ ratio (CLR_2_) transformation to identify relative compositional shifts, with CLR_2_-transformed values summing to zero within each sample. After adjustment for sample set, AD was associated with higher relative abundance of PC O-, lyso-phosphatidate (LPA), hexosylceramide (HexCer), and SM species and lower relative abundance of PE, lyso-phosphatidylcholine (LPC), lyso-phosphatidylethanolamine (LPE), lyso-phosphatidylserine (LPS), and lyso-phosphatidylinositol (LPI) species, reaching significance at the lipid-class level (**Figure 1c**). A significant trend toward more negative CLR_2_ values emerged in the AD brain with increasing lipid acyl chain unsaturation (**Figure 1d**), driven primarily by PE O- and PS species (**Extended Data Figure 3a**), with docosahexaenoic acid (DHA)-containing lipids showing uniform relative depletion across lipid classes. Interestingly, SMs trended in the opposite direction, with more very-long-chain SMs with a higher degree of unsaturation in the AD brain (**Extended Data Figure 3a**). Comparison of brain lipid class-level CLR_2_ differences for *APOE3/4* vs. *APOE3/3* and AD vs. control suggests that the *APOE4*-associated compositional shift broadly paralleled that of AD (**Figure 1e**, **Extended Data Figure 3b**). Sex additionally modified brain lipidomic profiles (**Extended Data Figure 3c**).

Lipid abundance analysis in human CSF revealed a striking trend of broad lipid depletion in AD, with a majority of lipid classes significantly decreased (**Figure 1f**). Hierarchical clustering of z-score normalized CSF lipid abundances showed the greatest separation by AD status, with *APOE3/3* controls exhibiting the highest CSF lipid abundances (**Extended Data Figure 2d**). More highly unsaturated lipids were relatively depleted in AD CSF (**Figure 1g**), similarly to but less pronounced than in the brain. In contrast, Cer and DG classes were relatively enriched in AD CSF (**Extended Data Figure 3d**). *APOE4* status and female sex produced comparatively modest shifts in CSF lipid composition by CLR_2_ analysis (**Extended Data Figure 3e–f**), but sex impacted both extent (stronger in females) and types of lipids reduced in AD (**Extended Data Figure 3g–h**). In serum, lipid profiles lacked comparably strong separation by AD or *APOE4* status (**Extended Data Figure 2e**, **Extended Data Figure 3i–j**), although HexCer was significantly decreased in *APOE4* carriers and sex-dependent differences were observed (**Extended Data Figure 3k**), reinforcing the compartment-specific nature of the CSF signature.

We next sought to understand to what extent AD-associated CSF lipid changes mirrored those of *APOE4*. Comparison of CSF lipid class-level log_2_ fold-changes for *APOE3/4* vs. *APOE3/3* against AD vs. control revealed largely concordant trends except for myelin-associated lipid classes (HexCer and sulfatides, Sulf), which displayed distinct behavior in *APOE4* carriers compared to individuals with AD (**Figure 1h**). Regression analysis underscored the broad decrease in CSF lipids associated with AD status, with significantly depleted lipid classes including PC, PC O-, PE, PG, LPC, LPG, TG, HexCer, SM, and Sulf (**Figure 1i**). *APOE4* status and female sex alone did not significantly alter the abundance of any lipid classes in CSF.

The most abundant lipids identified in the CSF included cholesterol (noting the logarithmic y-scale), PC 16:0-18:1 (POPC), cholesteryl esters (CEs 18:2, 18:1, 16:0, 20:4, with enriched saturated/mono-unsaturated fatty acids relative to serum as recently demonstrated;^25^ **Extended Data Figure 3l**), and various other phospholipids (**Figure 1j**). Importantly, these lipid classes are established components of CSF lipoprotein particles (LPs) that facilitate lipid transport in the brain (**Figure 1k**),^26–29^ thus the observed CSF lipid depletion may reflect reduced CSF LPs.

We then aimed to connect these prominent lipid changes to cognition. As anticipated, global cognitive function was strongly, inversely correlated with amyloid burden in the brain. All ten most abundant CSF lipids were positively correlated with global cognitive function, with two significantly correlated PC species (**Figure 1l**). Overall, these findings support disrupted lipid mobilization and transport in AD, characterized by distinct lipid remodeling across the brain and reduced CSF lipid abundance associated with poorer cognitive function.

### Lipidomic profiling of humanized *APOE* mice

As our human cohort consisted entirely of older donors, we could not determine the effect of age on lipidomic profiles. To investigate how age, *APOE* genotype, and chromosomal sex shape lipid profiles, we profiled a humanized *APOE* mouse cohort (n ≥ 6 each for combinations of young, 3 months vs. aged, 12 months; humanized *APOE2/2, APOE3/3,* or *APOE4/4* genotype; and female vs. male; **Figure 2a**). Matched brain, CSF, and blood plasma were collected from each mouse and analyzed by mass spectrometry-based lipidomics, identifying 239 unique, annotated lipids within 6,143 total detected lipid features after quality-control filtering. Similarly to the human cohort, lipid profiles correlated most strongly within a given compartment (**Extended Data Figure 4a**) and formed clear tissue/biofluid-specific clusters by PCA (**Extended Data Figure 4b**).

**Figure 2.**
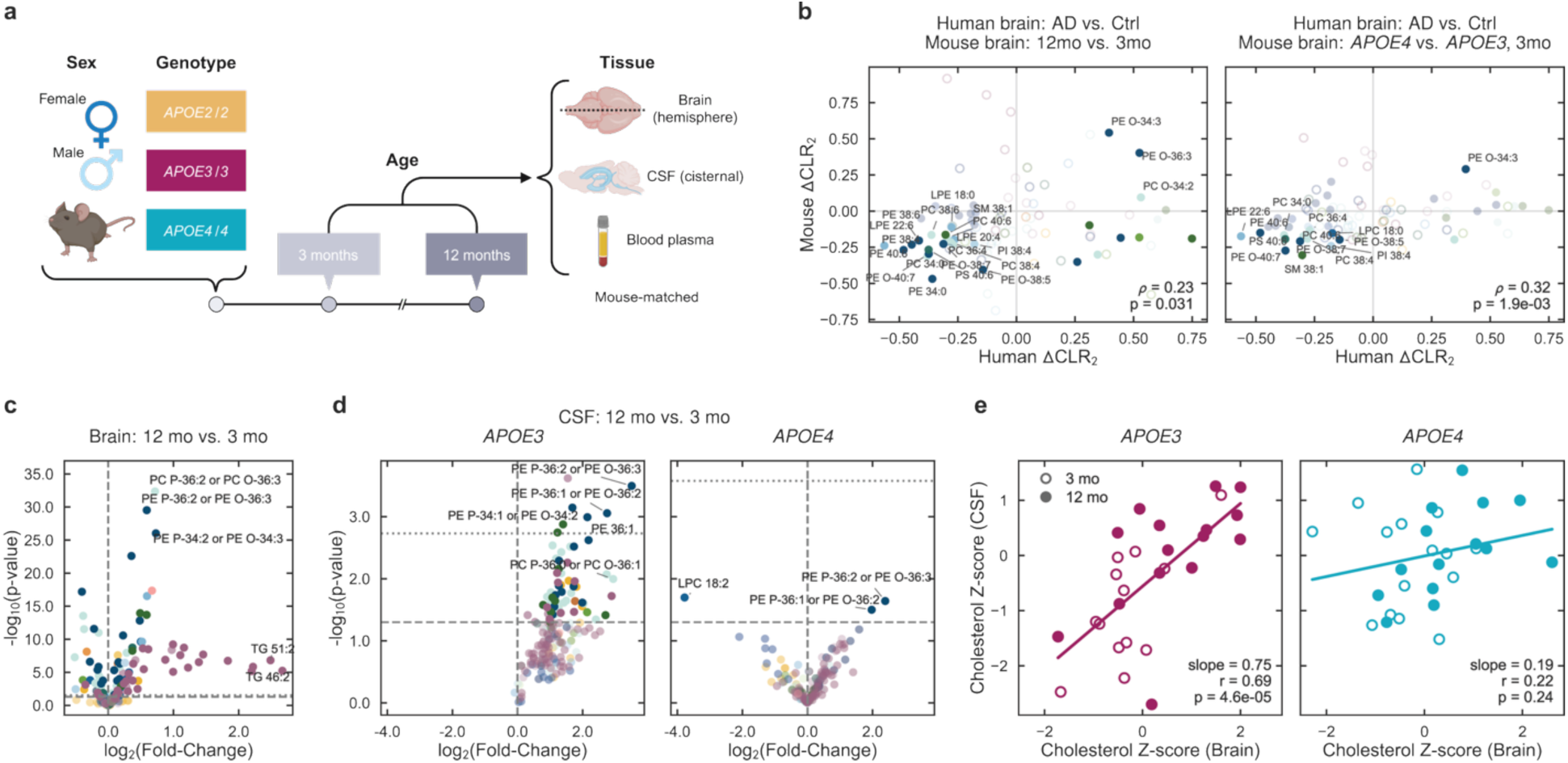
Aging drives brain lipid remodeling and *APOE*-dependent CSF lipid responses in mice. **(a)** Overview of mouse study design to compare lipid changes in young (3 months) vs. aged (12 months) mice with humanized homozygous *APOE2*, *APOE3*, or *APOE4* genotypes, for female and male mice. Brain tissue, cisternal CSF, and blood plasma matched per mouse were analyzed by mass spectrometry-based lipidomics. **(b)** Mouse brain lipid CLR_2_ difference between aged vs. young mice including all genotypes (left) and between *APOE4* vs. *APOE3* in young mice (right) against human brain lipid CLR_2_ difference between AD vs. Ctrl. Acyl chain-resolved lipids in the human dataset were summed across sidechains to determine matched lipids in the mouse dataset. Markers indicate individual lipid species colored by lipid class, with filled markers for *P* < 0.05 in human ΔCLR_2_ and shaded fully opaque for *P* < 0.05 in both species. Annotation provides Spearman’s correlation coefficient *ρ* and corresponding *P* value. Volcano plots (-log_10_ *P* value vs. log_2_ fold-change) of **(c)** brain lipids including all genotypes and **(d)** CSF lipids for aged vs. young mice for *APOE3* (left) and *APOE4* (right). Markers indicate lipid species, grouped by color for different lipid classes. *P* values were determined using a two-sided *t*-test assuming equal variances. Short-dashed line indicates FDR-adjusted *P* = 0.05 (Benjamini-Hochberg correction). **(e)** Z-score normalized cholesterol abundance in CSF vs. brain, separated by *APOE3* (left) and *APOE4* (right). Marker fill denotes age (3 months: open, 12 months: filled). Lines indicate ordinary least-squares regression fits; *r* values are shown, with slope significance determined by a two-sided t-test.

Lipidomic changes observed in the human AD vs. control brain were modestly concordant with those in mouse aging as well as *APOE4* vs. *APOE3* comparisons (**Figure 2b**). Within the mouse brain, age was the dominant axis of lipidomic variation (**Extended Data Figure 4c**), characterized predominantly by increased lipid abundance (**Figure 2c**). Specific PC, PE, TG, and other species increased significantly, with broad accumulation of TGs consistent with increased neutral-lipid storage with age. Age-associated TG accumulation was compositionally selective, with the greatest increases among short-carbon-chain, low-to-moderately unsaturated TG species, while sparing both short-chain, fully saturated TGs and long-chain, highly polyunsaturated TGs (**Extended Data Figure 4d**). As TGs within tissue are predominantly stored in lipid droplets, these findings suggest neutral-lipid storage programs with compositional windows set by both total acyl chain length and degree of unsaturation. These age-associated changes were also sex dependent: Aged female brains showed elevated TGs and reduced CEs relative to males (**Extended Data Figure 4e**), despite increased cholesterol with aging across both sexes.

In CSF, a pronounced *APOE*-dependent response to aging emerged: Lipid abundance broadly increased with age in *APOE3/3* mice, but not in *APOE4/4* (**Figure 2d**). This pattern was especially evident among CEs, a major LP-associated class, with significant increases in longer carbon chain, more unsaturated CEs with age (**Extended Data Figure 4f**). Comparison of brain and CSF lipid profiles demonstrated increased brain lipid abundance with age across all three genotypes, but a commensurate increase in CSF lipids only in *APOE3/3* and *APOE2/2* mice (**Extended Data Figure 4g**). These data thus reveal an age-adaptive increase in CSF lipids in *APOE3/3* and *APOE2/2* mice that is absent in *APOE4/4* mice.

Cholesterol, as the most abundant brain and CSF lipid, further illustrated this genotype-dependent coupling between brain and CSF: Higher brain cholesterol was significantly correlated with higher CSF cholesterol in *APOE3/3* mice, whereas *APOE4/4* mice did not demonstrate this relationship (**Figure 2e**). Extending this analysis across all lipids, most brain lipids in *APOE3/3* mice showed strong positive correlations with CSF lipid abundances, whereas the relationship was weaker in the *APOE4/4* mouse brain and CSF (**Extended Data Figure 4h**). In contrast to these positive correlations between brain and CSF, many TG, DG, and PE lipids in plasma were negatively correlated with CSF lipid abundances in *APOE3/3* mice, while *APOE4/4* mice exhibited weak or even positive correlations. In sum, the mouse lipidomic analysis indicates that aging drives brain lipid remodeling, which was modified by *APOE* genotype and sex, producing a lipid signature partially recapitulating that of the human AD brain. In CSF, *APOE4* mice did not exhibit age-related increases in LP-associated lipids, paralleling the reduced lipid abundance observed in human AD CSF and suggesting disrupted lipid mobilization and transport in aged *APOE4* mice.

### Engineered rLPs facilitate biomolecule binding and lipid efflux

Based on these findings, we hypothesized that supplementation of the CSF with functional LPs could rescue lipid-transport defects arising in the AD and *APOE4* brain. Brain-derived LPs are ∼10–30 nm diameter endogenous nanoparticles composed of phospholipids, cholesterol, and apolipoproteins – predominantly ApoE, as well as ApoA-I and ApoJ.^26–28^ Nascent ApoE-containing LPs are secreted primarily by astrocytes as discoidal particles (**Figure 3a**) and can undergo maturation into spherical LPs with CE-rich cores as found in the CSF.^1,27^ Guided by our human CSF lipidomics data together with synthetic feasibility considerations, we engineered brain-inspired reconstituted lipoprotein particles (rLPs) containing the non-AD-risk ApoE3 protein and abundant yet stable CSF lipids. Rather than recreate mature, spherical CSF LPs, we purposely generated nascent, discoidal rLPs to promote rLP capacity to accept cellular lipids. Specifically, we included the two most abundant lipids, cholesterol and PC 16:0–18:1 (POPC) (**Figure 1j**), as well as two highly abundant PEs, a lipid class particularly enriched in the brain and important for membrane curvature. Based on the criterion of no more than one double bond to limit susceptibility to oxidation, we selected PE 18:0–18:1 (SOPE) and PE 16:0–18:1 (POPE) (**Figure 1j**). Importantly, CSF abundances of both POPC (**Figure 1l**) and SOPE were significantly associated with the individual’s global cognitive function.

rLPs were synthesized by thin-film hydration followed by cholate-mediated ApoE incorporation and disc self-assembly, then purified by size-exclusion chromatography (SEC) (**Extended Data Figure 5a**; full details in Methods). From SEC, ApoE and the lipophilic dye (Vybrant DiI) co-eluted based on overlapping absorbance maxima at 280 nm and 550 nm, respectively (**Figure 3b**), supporting formation of ApoE-associated nanoparticles. Dynamic light scattering (DLS) revealed a unimodal population with a mean number-weighted hydrodynamic diameter of 23.5 nm, distinct from non-lipidated ApoE at 5.6 nm (**Figure 3c**). rLPs also exhibited a more negative zeta potential than non-lipidated ApoE, −39.1 mV vs. −11.1 mV, respectively, suggesting enhanced colloidal stability upon lipidation (**Figure 3d**). Transmission electron microscopy (TEM, **Figure 3e**) and cryogenic TEM (**Extended Data Figure 5b**) further revealed nanoscale discoidal particles consistent with nascent brain LP morphology. rLPs were observed in both face-on and edge-on orientations, with average diameter of 22.9 ± 3.6 nm (consistent with those measured by DLS in the hydrated state) and thickness of 4.9 ± 1.0 nm (**Extended Data Figure 5c**), in line with a typical phospholipid bilayer thickness of ∼4–5 nm. In the absence of ApoE, particles with bending bicelle-like morphology were formed (**Extended Data Figure 5d**).

**Figure 3.**
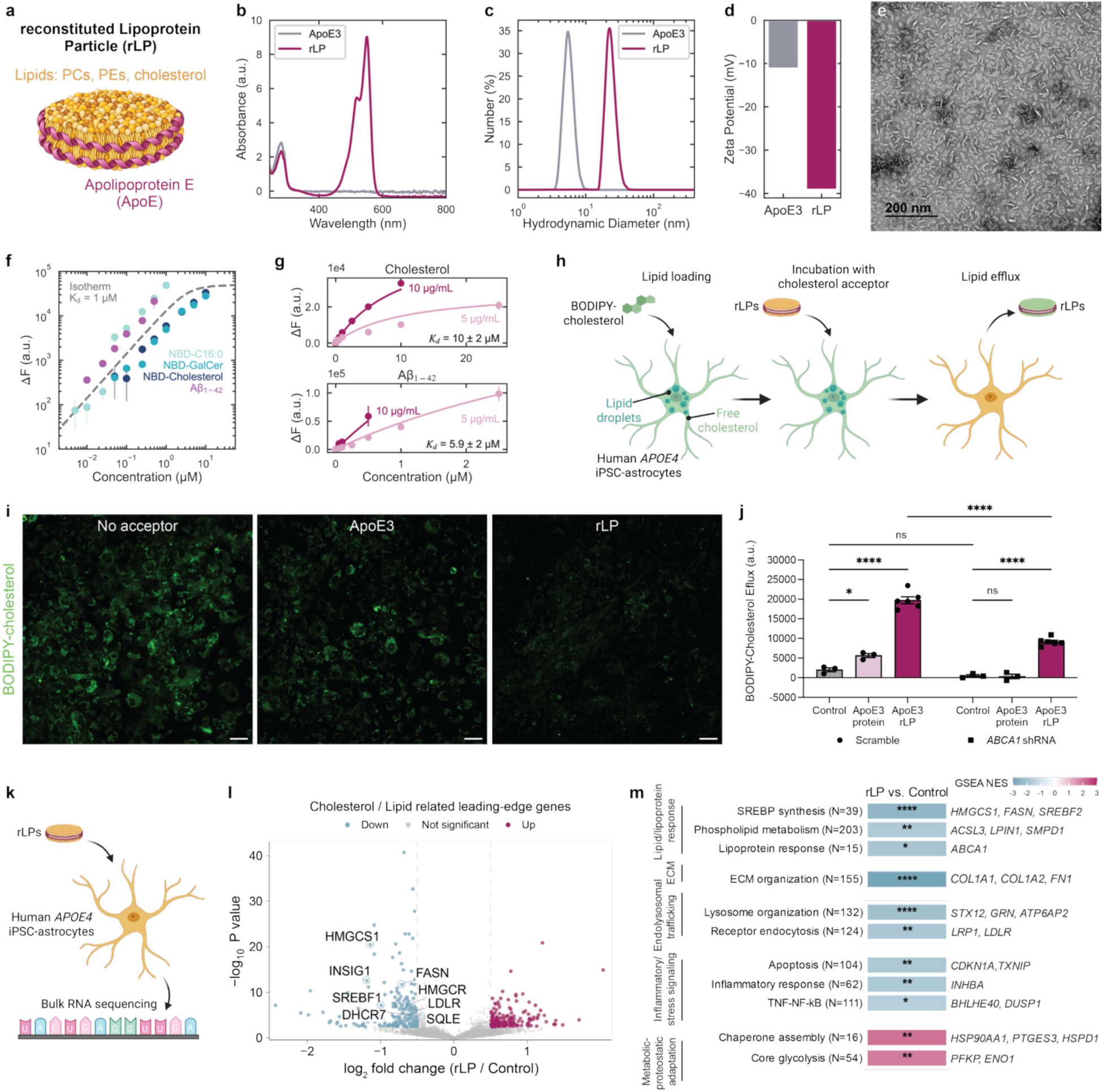
Brain-inspired reconstituted lipoprotein particles (rLPs) mediate biomolecule binding in solution and lipid efflux from human *APOE4* astrocytes *in vitro*. (a) Rendering of discoidal rLPs composed of lipids including PCs, PEs, and cholesterol (yellow), with ApoE (magenta). **(b)** Absorbance spectra of rLPs (magenta), with peaks at 280 nm for protein and 550 nm for the Vybrant DiI lipophilic dye, compared to free ApoE3 (gray). **(c)** Hydrodynamic diameter of rLPs (23.47 nm) compared to free ApoE3 (5.64 nm), measured by dynamic light scattering and reported as number distributions with mean number-weighted diameters. **(d)** Zeta potentials of rLPs (-39.09 mV) compared to free ApoE3 (-11.12 mV). **(e)** Discoidal morphology of rLPs visualized by transmission electron microscopy (TEM). **(f–g)** In-solution binding assays of nitrobenzoxadiazole (NBD)-labelled palmitic acid (C16:0), galactosylceramide (GalCer), and cholesterol, or HiLyte Fluor 647-tagged amyloid-β peptide 1–42 (Aβ42), incubated at varying concentrations with rLPs. NBD-lipid fluorescence was measured at 544 nm (excitation, 473 nm) and HiLyte-Aβ42 fluorescence was measured at 674 nm (excitation, 600 nm) following removal of unbound peptide. Fluorescence changes were calculated relative to corresponding lipid- or Aβ42-only controls (no rLPs). **(f)** Concentration-dependent fluorescence responses with 10 µg/mL rLPs, with the dashed line representing an illustrative binding isotherm (K_D_ = 1 µM). Error bars were calculated by propagating the standard errors of controls and replicates (*n* = 3). **(g)** Fluorescence responses for cholesterol (top) and Aβ42 (bottom) with 5 or 10 µg/mL rLPs. Points represent mean ± SEM (*n* = 3); solid lines indicate fitted binding isotherms, with apparent dissociation constants (K_D_) and fitting standard errors indicated. **(h)** Schematic of cholesterol-efflux assay in which human induced pluripotent stem cell (iPSC)-derived *APOE4* astrocytes were loaded with BODIPY-tagged cholesterol, treated with rLPs for 24 h, and assessed for BODIPY signal in cells and medium. **(i)** Representative images of BODIPY-cholesterol-loaded *APOE4* astrocytes after treatment with no acceptor, free ApoE3, and rLPs. Scale bars, 50 µm. **(j)** Quantification of BODIPY-cholesterol fluorescence signifying lipid efflux into the medium for *APOE4* astrocytes pre-treated with scramble (left) or *ABCA1* (right) shRNA then treated with no acceptor, free ApoE3, or rLPs. Markers denote individual wells (averaged over *n* = 4 regions per well). Error bars show mean ± SEM (*n* ≥ 3). Statistical significance was determined using unpaired two-sided t-test. * *P* < 0.05, **** *P* < 0.0001. **(k)** Schematic of *in vitro* rLP treatment and bulk RNA-sequencing of *APOE4* astrocytes. **(l)** Volcano plot of the astrocyte transcriptional response to rLP treatment measured by bulk RNA sequencing (*n* = 5 per group), with selected GSEA leading-edge genes from cholesterol- and lipid-related pathways highlighted. Colors indicate genes meeting FDR < 0.05 and |log_2_ fold-change| > 0.5. **(m)** Gene-set enrichment analysis (GSEA) of *APOE4* astrocytes treated with rLPs vs. control. Colors indicate normalized enrichment scores (NES), with blue and pink denoting negative and positive enrichment, respectively. Representative directionally concordant genes passing gene-level FDR < 0.05 are shown at right. *N* indicates the number of genes in each gene set. GSEA significance: * FDR < 0.05, ** FDR < 0.01, *** FDR < 0.001, **** FDR < 0.0001.

To assess functionality, we first examined the capacity of rLPs to acquire exogenous lipid cargo characteristic of the brain. We employed a polarity sensitive fluorophore, nitrobenzoxadiazole (NBD), to track transfer from the free, aqueous state (quenched, red-shifted fluorescence) into the hydrophobic nanoparticle environment (increased intensity, blue-shifted fluorescence). NBD-labeled palmitic acid (C16:0), galactosylceramide (GalCer), and cholesterol each exhibited concentration-dependent increased fluorescence in the presence of rLPs (**Extended Data Figure 5e**, **Figure 3f**). Given the known interaction between ApoE and amyloid-β (Aβ),^9,31^ we next asked if this binding capacity extended beyond lipids. We found that fluorescently labeled amyloid-β peptide 1–42 (Aβ42) was enriched in the rLP-containing fraction following size-dependent separation of unbound peptide, supporting an association between rLPs and Aβ42 that could be leveraged for additional therapeutic effect (**Extended Data Figure 5e**, **Figure 3f**). By fitting the concentration-dependent fluorescence responses of lipid or Aβ42 to a site-binding isotherm, we estimate apparent dissociation constants (*K*_D_) of 10 ± 2 µM for cholesterol, 9.2 ± 1 µM for GalCer, 0.52 ± 0.1 µM for C16:0, and 5.9 ± 2 µM for Aβ42 (**Figure 3g**, **Extended Data Figure 5f**). These findings demonstrate the capacity of rLPs to associate with diverse lipid species and Aβ42, supporting their potential as multifunctional biomolecular carriers.

We next tested if this lipid-acceptor capacity could promote cellular lipid mobilization. Human *APOE4* iPSC-derived astrocytes were loaded with BODIPY-labelled cholesterol, followed by rLP treatment and quantification of both cellular lipid droplets (LDs) and cholesterol detectable in the medium (**Figure 3h**). rLP treatment reduced astrocyte LD burden by 78.1 ± 2.3%, while concurrently increasing BODIPY-cholesterol in the medium by 5.4 ± 0.2-fold (mean ± standard error of the mean; **Figure 3i–j**, **Extended Data Figure 6a–b**). Neither free ApoE3, ApoE4, ApoA-I, nor lipoprotein particles lacking ApoE3 promoted cholesterol efflux to the same extent as rLPs. Perturbation of canonical cholesterol-efflux pathways^9^ attenuated this response: Pharmacological inhibition of ABCA1-dependent cholesterol efflux with probucol^32^ or LXR signaling with GSK2033^33^ partially blunted the rLP-mediated reduction in lipid burden by 31.0 ± 8.4% and 60.8 ± 11.5%, respectively (**Extended Data Figure 6c–e**). To more directly test the contribution of these canonical cholesterol-efflux pathways, shRNA-mediated knockdown of *ABCA1* (**Extended Data Figure 6f**) similarly decreased rLP-mediated cholesterol efflux by 53.5 ± 2.9% (**Figure 3j**), whereas *ABCG1* knockdown reduced it by only 12.8 ± 3.5% (**Extended Data Figure 6g–h**). In comparison, *ABCA1* knockdown led to nearly complete blocking of cholesterol efflux by free ApoE3 (94.2 ± 10.4%; **Figure 3j**), suggesting that ABCA1-independent mechanisms also contribute to rLP-mediated cholesterol efflux from astrocytes.

The effect of rLP treatment extended beyond supporting cholesterol efflux. Bulk RNA-sequencing of *APOE4* astrocytes following rLP treatment (**Figure 3k**) revealed coordinated transcriptional changes across pathways involved in lipid synthesis, uptake, intracellular trafficking, inflammation, and cellular stress (**Figure 3l–m**). In particular, the SREBP-regulated lipid synthesis pathway was negatively enriched, with reduced expression of multiple cholesterol-and lipid-synthesis genes, including *SREBF1, SREBF2*, *HMGCS1*, and *FASN*, consistent with reduced compensatory cholesterol biosynthesis associated with disrupted lipid homeostasis in *APOE4* astrocytes.^17,34^ Pathways involved in lipoprotein uptake and receptor-mediated endocytosis were also negatively enriched, including reduced expression of *LDLR* and *LRP1*, as were lysosomal organization and endosomal trafficking pathways. These findings may reflect changes in intracellular lipid processing following rLP-mediated cholesterol mobilization. Beyond lipid metabolism, rLP treatment was also associated with negative enrichment of ECM organization pathways, with reduced expression of *COL1A1*, *COL1A2*, and *FN1*, consistent with attenuation of astrocytic matrisome dysregulation previously reported in *APOE4* astrocytes.^34,35^ Inflammatory and stress-related pathways were also negatively enriched, including TNF–NF-κB signaling and apoptosis, whereas chaperone assembly and glycolysis were positively enriched. Collectively, these findings support that ApoE3-rLP-mediated cholesterol efflux is accompanied by a coordinated reduction in compensatory lipid synthesis, uptake, and intracellular processing, potentially alleviating the dysregulated lipid homeostasis in *APOE4* astrocytes.

### rLP treatment reduces lipid and amyloid-β burden *in vivo*

We translated rLP treatment *in vivo* to test if this approach modified Alzheimer’s-associated pathology within the full brain context. rLPs or saline control were administered by intracisternal injection to 12–14-month-old humanized *APOE4* mice crossed with the 5xFAD Alzheimer’s model (*APOE4*;5xFAD) (**Figure 4a**; full details in Methods). By 24 h, rLP treatment significantly reduced lipid droplet accumulation in the subiculum (**Figure 4b–c**) and Aβ burden in the hippocampus (**Figure 4d–e**), as visualized in the dentate gyrus (DG) and CA1 regions and quantified by both total Aβ coverage and plaque count (**Extended Data Figure 7a**). Across matched animals, lipid droplet and Aβ burden tracked together (**Figure 4f**). At 12–14-month age, lipid droplet and Aβ content were not significantly associated with mouse sex, parental origin of the 5xFAD transgene, or exact age, except for the expected correlation between Aβ and age (**Extended Data Figure 7b**). Significant reductions in Aβ coverage were also observed in the prefrontal cortex (**Extended Data Figure 7c–d**). Interestingly, rLP treatment led to similar changes – enriched TGs and depleted CEs – in *APOE4;*5xFAD mouse brains as were previously seen with age in *APOE3/3* relative to *APOE4/4* mouse brains (Pearson correlation coefficient *r* = 0.63; **Extended Data Figure 7e**).

**Figure 4.**
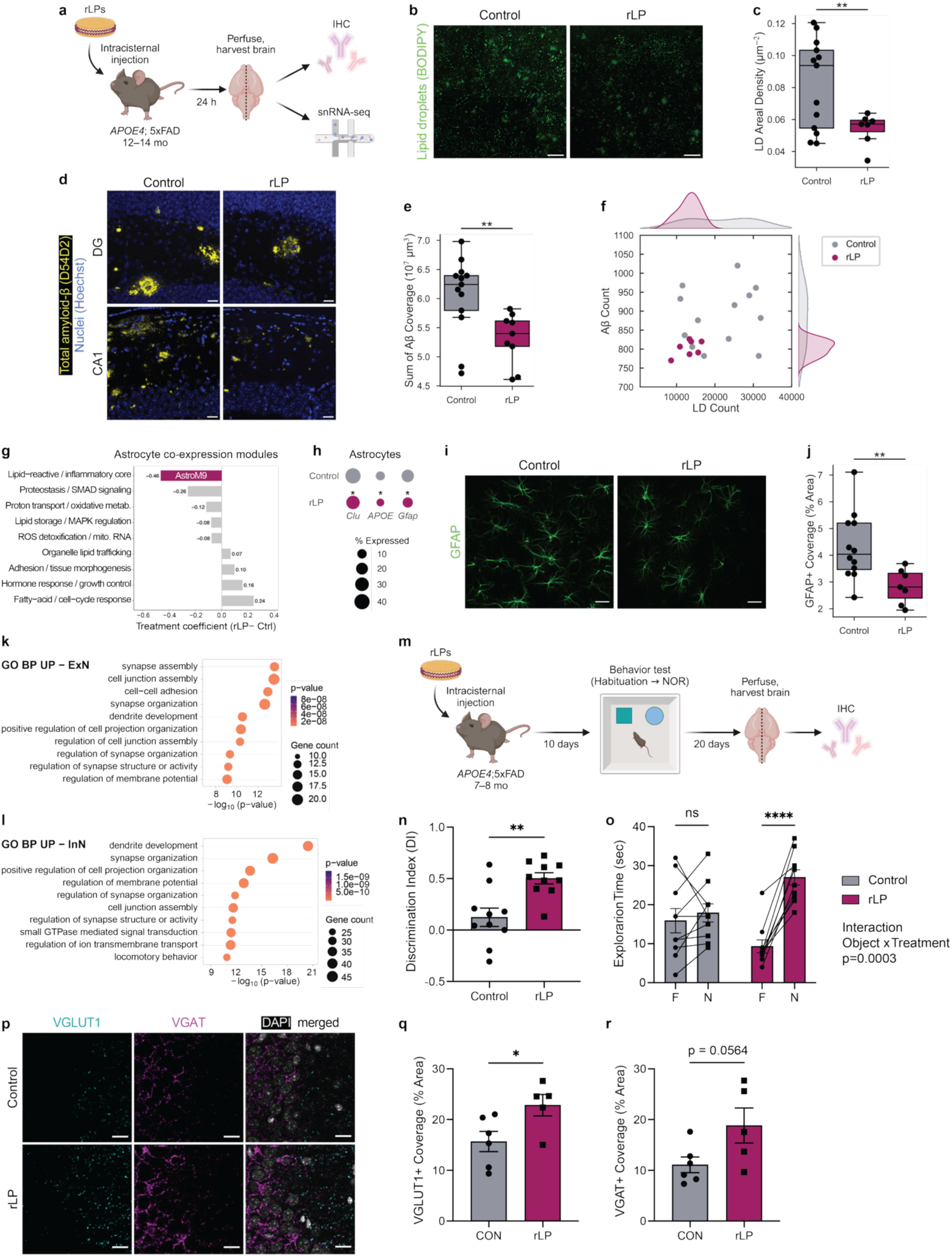
rLP treatment reduces lipid and amyloid-β burden, promotes synaptic remodeling, and improves novel object recognition in *APOE4*;5xFAD mice. (a) 12–14-month-old *APOE4;*5xFAD mice were intracisternally injected with saline control or rLPs and euthanized after 24 h. **(b)** Representative images of BODIPY-stained lipid droplets (LDs; green) in mouse subiculum for control (left) and rLP treatment (right), with corresponding **(c)** quantification of LD count normalized to subiculum area for control (gray) vs. rLP treatment (magenta) (control *n* = 13, rLP *n* = 7). Scale bars, 50 µm. **(d)** Representative images of D54D2-stained total Aβ (yellow) and Hoechst-stained cell nuclei (blue) in mouse hippocampus for control (left) and rLP treatment (right) in the dentate gyrus (DG; top) and CA1 (bottom). Scale bars, 20 µm. **(e)** Quantification of net hippocampal Aβ coverage for control vs. rLP treatment (control *n* = 12, rLP *n* = 9). **(c,e)** Markers denote individual mice and statistical significance was determined using Welch’s two-sided t-test. **(f)** Aβ count in the hippocampus vs. LD count in the subiculum, paired within the same mouse. Corresponding PDFs by KDE are included on the top and side for LDs and Aβ, respectively. **(g)** Adjusted treatment effects on astrocyte co-expression modules (control *n* = 7, rLP *n* = 9), controlling for sex and parental 5xFAD origin. Negative coefficients indicate reduced module scores following rLP treatment. AstroM9 is highlighted in magenta. **(h)** Key DEGs in astrocytes. **(i)** Representative images of GFAP staining (green) in mouse prefrontal cortex for control (left) and rLP treatment (right) with corresponding **(j)** quantification of GFAP-positive cell coverage as a percent area (control *n* = 12, rLP *n* = 7). Scale bars, 200 µm. **(k-l)** GO enrichment analysis of top 10 pathways in **(k)** excitatory neurons (ExN) and **(l)** inhibitory neurons (InN). **(m)** 7–8-month-old *APOE4;*5xFAD mice were intracisternally injected with control or rLPs (control *n* = 10, rLP *n* = 10). Mice recovered for one week before undergoing the novel object recognition test after habituation to the behavioral room and arena. **(n)** Discrimination index (DI) for control and rLP treatment groups. Positive DI values indicate preferential exploration of the novel object (see Methods). **(o)** Exploration time for the familiar (F) and novel (N) objects for each animal. Each data point and connecting line indicate an individual animal. Two-way ANOVA revealed a significant object × treatment interaction (*P* = 0.0003). **(p)** Representative images of VGLUT1 (cyan) and VGAT (magenta) immunostaining in hippocampal CA3 of control (left) and rLP treatment (right). Scale bars, 10 µm. Quantification of **(q)** VGLUT1- and **(r)** VGAT-positive coverage as percent area (control *n* = 6, rLP n = 5). **(n, o, q, r)** Data are mean ± SEM; markers denote individual mice. Two-tailed unpaired t-test. * *P* < 0.05, ** *P* < 0.01, *** *P* < 0.001, **** *P* < 0.0001.

To capture the broader cellular transcriptomic response to rLP treatment, we performed single-nucleus RNA sequencing (snRNA-seq) of hippocampi from the same mice. After quality-control and doublet filtering, we recovered 300,399 nuclei spanning eight major cell populations, including excitatory and inhibitory neurons, astrocytes, choroid plexus epithelial cells, microglia, oligodendrocytes, oligodendrocyte precursor cells, and vascular cells comprising endothelial cells, pericytes, vascular smooth muscle cells, and fibroblasts (**Extended Data Figure 8a–b**). No significant differences in cell-type proportions were detected between treatment groups (**Extended Data Figure 8c–d**). rLP treatment resulted in 1,028 upregulated and 544 downregulated genes, with inhibitory neurons displaying the largest transcriptional response (**Extended Data Figure 8e**).

Astrocytes from rLP-treated mice showed reduced expression of AstroM9, a co-expression module enriched for lipid-reactive and inflammatory genes (**Figure 4g**). Consistent with findings from rLP-treated *APOE4* iPSC-derived astrocytes, both models showed downregulation of pathways related to lipid storage and lysosomal processing, inflammatory responses, and extracellular-matrix remodeling. These convergent changes suggest that rLP treatment may alleviate lipid-handling stress in *APOE4* astrocytes and reduce the requirement for compensatory cholesterol biosynthesis. Within AstroM9, *APOE*, *CLU*, and *GFAP* were downregulated (**Figure 4h**). Reduced endogenous *APOE4* expression raises the possibility that supplementation with exogenous ApoE3-containing rLPs influences astrocytic ApoE4 production, whereas reduced *GFAP* expression is consistent with attenuation of a reactive astrocyte state. Corroborating these findings at the protein level, we observed reduced GFAP-positive cell coverage in the prefrontal cortex of rLP-treated mice (**Figure 4i–j**).

Gene Ontology (GO) analysis of genes upregulated in excitatory and inhibitory neurons revealed enrichment of pathways associated with synapse assembly and organization, dendrite development, neuronal connectivity and maturation, and regulation of membrane excitability (**Figure 4k–l**). AstroM9 scores were inversely associated with broad neuronal-expression programs, particularly inhibitory-neuron programs (**Extended Data Figure 9a–c**). Thus, the reduction in AstroM9 following rLP treatment may reflect an astrocytic state associated with improved support of neuronal function, although these correlations do not establish causality. Within the neuronal programs, both neuronal populations upregulated genes involved in axon guidance and neurite organization (*Dcc, Robo1, Efna5, Dscam, Cobl*) and neurotransmission and excitability (*Grid1, Grid2*, *Asic2*), alongside cell-type-specific increases in genes related to synaptic organization, neurite structure and excitability in inhibitory neurons (*Nlgn1, Shank3, Gap43, Slit1, Map2, Cacna1e*) and neuronal development and plasticity in excitatory neurons (*Ntrk2, Reln, Adcy8, Mef2c*) (**Extended Data Figure 9e**).

Microglia also showed downregulation of lipid-stress, disease-associated microglial, and phagolysosomal programs, consistent with attenuation of lipid-processing and lysosomal stress in glial cells (**Extended Data Figure 9e–f**), although there were no significant changes in IBA1-positive cell coverage in the rLP-treated group hippocampus or prefrontal cortex (**Extended Data Figure 9g**). Overall, these transcriptional changes indicate that the acute effects of rLP treatment extend beyond reduced lipid and Aβ burden to cell-type-specific responses associated with neuronal function and glial state.

### rLP treatment improves recognition memory in *APOE4*;5xFAD mice

Given our snRNA-seq findings of altered synaptic organization following rLP treatment and the role of ApoE-containing LPs in neuronal lipid support,^1,36^ we next asked whether rLP treatment could improve recognition memory as a behavioral measure of cognitive function. 5xFAD mice, a preclinical model of AD, begin to display cognitive impairment at approximately 4–5 months of age.^37^ Therefore, we injected rLPs into 7–8-month-old *APOE4*;5xFAD mice, an age at which cognitive impairment is expected but overt motor deficits are not prominent.^37^ Ten days after injection, we performed the novel object recognition test following habituation to the testing room and arena (**Figure 4m)**. During the test session, total distance travelled and total object exploration time did not differ between groups **(Extended Data Figure 10a–b)**. *APOE4*;5xFAD mice displayed cognitive impairment, represented by discrimination index (DI) that is closer to zero, which indicates equal exploration of the novel and familiar objects, while a positive value indicates preferential exploration of the novel object (**Figure 4n**). Strikingly, rLP-treated *APOE4*;5xFAD mice showed a significantly higher DI than control mice (**Figure 4n**), spending significantly more time exploring the novel object than the familiar object, whereas control mice showed no significant preference (**Figure 4o**).

To determine whether improved recognition memory was accompanied by changes in synaptic composition, as suggested by our snRNA-seq data, we immunostained for the vesicular glutamate transporter VGLUT1 and the vesicular GABA transporter VGAT, markers of glutamatergic and GABAergic presynaptic terminals, respectively. Our quantification revealed that rLP treatment significantly increased VGLUT1-positive coverage (*P* = 0.0367) and elevated VGAT-positive coverage (*P* = 0.0564) in hippocampal CA3, compared to control mice **(Figure 4p–r)**. Taken together, these findings suggest that rLP treatment improves memory and promotes synaptic remodeling in the hippocampus of *APOE4*;5xFAD mice.

## Discussion

ApoE-LP-mediated lipid transport is critical in regulating the brain lipid economy, shaping lipid availability, distribution, and storage across local cellular and broader tissue-level scales. Our findings reveal altered brain and CSF lipid composition associated with AD and *APOE* genotype, together with cellular responses to ApoE-LP perturbations.

Across human brain and CSF, AD and *APOE4* were associated with related but compartmentally distinct lipidomic shifts. The AD brain showed relatively enriched complex sphingolipid classes, including SM and HexCer, as well as relative depletion of several phospholipid classes and of more highly unsaturated species, including lipids containing DHA, a neuroprotective omega-3 fatty acid. Prior human brain lipidomics studies have similarly reported enrichment of glycosphingolipids and longer-chain SM species in AD,^38^ increased HexCer alongside reduced lysophospholipids in *APOE4* carriers,^39^ and AD-associated depletion of PE, LPE, LPC, and phospholipids containing DHA.^40–43^ The similar brain lipidomic signature between AD and *APOE4* supports convergence between genetic risk and disease-related lipid remodeling.

In the CSF of individuals with AD, particularly among females, we additionally uncovered extensive depletion of LP-associated lipids, consistent with trends reported in independent lipidomic analyses of lumbar CSF.^29,44^ Our observation that LP-relevant CSF lipids tended to correlate positively with cognition also aligns with previous work linking higher CSF LP concentrations to better cognitive performance.^29^ Reduced CSF cholesterol-efflux capacity, consistent with impaired LP-mediated lipid transport, has also been reported in AD.^45^ Ceramides showed the greatest AD-associated increase in relative abundance among CSF lipid classes, consistent with previous reports of elevated ceramides in AD CSF,^46^ and enrichment of ceramides in amyloid-β plaques.^47,48^ While AD and *APOE4* were associated with similar lipidomic shifts in the brain, this correspondence was weaker in CSF, particularly for myelin-associated HexCer and Sulf classes. Our human lipidomic analyses extend prior work by resolving differential lipid abundances across matched brain, CSF, and serum and by stratifying effects across *APOE* genotype and sex.

In mice, aging led to brain lipid remodeling, characterized by general lipid accumulation in the brain that showed sex-dependent divergence in neutral lipid storage. These brain lipid changes were accompanied by changes in CSF lipid abundance. Notably, age-related increases in CSF cholesterol and cholesteryl esters were evident in *APOE3* and *APOE2* mice but absent in *APOE4* mice, suggesting that ApoE4 may limit adaptive lipid redistribution as brain lipid burden increases with age. More broadly, our human and mouse lipidomic datasets provide a resource for mapping lipid changes in AD and its associated risk triad, identifying candidate lipid biomarkers, and nominating potential targets for therapeutic intervention.

Motivated by these observations, we hypothesized that supplying functional ApoE3-containing lipoprotein particles could restore lipid transport capacity. We leveraged our CSF lipidomic profiling to rationally design and synthesize brain-inspired discoidal rLPs, mimicking the morphology of nascent lipoproteins to facilitate lipid uptake.^1,27^ We found that rLPs acted as lipid acceptors in promoting cholesterol efflux from human iPSC-derived *APOE4* astrocytes through canonical cholesterol-export pathways, including ABCA1, and induced coordinated transcriptional remodeling spanning lipid synthesis, uptake, intracellular trafficking, inflammatory signaling, and cellular metabolism. Moreover, the ability of rLPs to associate with Aβ and to improve lipid, Aβ, and neuronal phenotypes *in vivo* raises the possibility that restoring ApoE-mediated lipid transport could influence multiple interconnected features of AD pathology. Prior work has established that glia-derived cholesterol carried by ApoE-containing LPs promotes synapse formation,^36^ and ApoE3-LPs enhance neurite outgrowth compared with ApoE4-LPs.^49^ More recently, CSF LP-mediated cholesterol delivery to neurons *in vitro* was found to be impaired in AD, with ApoE4-LPs showing reduced neuronal cholesterol delivery compared with ApoE3-LPs.^50^ Our findings also contextualize prior work that HDL-like particles can interact with Aβ and facilitate removal to improve cognition in mice.^51^

These findings are conceptually aligned with recent work showing that peripherally derived ApoA-I crosses the blood-brain barrier and engages astrocytic ABCA1 to drive cholesterol efflux, supporting synaptic integrity in 5xFAD mice.^52^ Together, these studies support the broader principle that restoring LP-mediated lipid homeostasis can improve neuronal function. By directly supplying functional, lipidated ApoE, our rLPs improve lipid, neuronal, and cognitive phenotypes in an *APOE4*-expressing AD model. Recent studies showing that ApoE-LPs can either promote neuronal lipid toxicity or remove harmful lipids highlight the importance of LP composition and the potential of our tunable rLP platform.^53,54^ Future work should clarify the mechanisms of how rLP-mediated treatment enhances Aβ clearance through CSF drainage pathways^55^ or cellular uptake and degradation,^56^ and how these mechanisms relate to lipid efflux and neuroprotection outcomes. More broadly, this work supports restoring brain lipid transport as a potential therapeutic avenue in AD, with engineered rLP nanoparticles offering a modular platform to investigate brain lipid transport and advance this approach.

## Materials and Methods

### Human cohort design and sample collection from the Rush Memory and Aging Project (MAP)

Human samples (prefrontal cortex, ventricular CSF, and blood serum) were sourced from donors in the MAP study, a longitudinal, epidemiologic clinical-pathologic cohort study of aging and dementia in elderly individuals from the greater Chicago metropolitan area.^24^ The study includes annual detailed clinical evaluation, postmortem pathological evaluations, and genetic, epigenomic, transcriptomic, proteomic, and metabolomic bulk-tissue profiling. All participants enrolled without known dementia. Informed consent and Anatomical Gift Act consent were obtained from each participant. The MAP Project was approved by an Institutional Review Board (IRB) of Rush University Medical Center. All participants signed a repository consent that allows their data and biospecimens to be shared.

We selected 68 individuals from the MAP study for which we obtained matching brain tissue, CSF, and, for the 40 individuals of the primary set, blood serum samples. Prefrontal cortex tissue (Brodmann area 10) was chosen because this region is highly implicated in the cognitive and behavioral changes affected by AD. Donors were chosen in the age range of 72–100 years and with a postmortem interval (PMI) ≤ 10 h, with no significant differences between AD vs. Ctrl and *APOE3/3* vs. *APOE3/4* subgroups, determined by the Mann-Whitney-Wilcoxon two-sided test (**Extended Data Figure 1a–b**). AD status was based on the National Institute on Aging (NIA)-Reagan Diagnosis criteria, which incorporates quantitative measures of pathology for both neurofibrillary pathology (Braak stage) and neuritic plaques (CERAD score) (**Extended Data Figure 1c**).^57,58^ A pathologic diagnosis of AD by NIA-Reagan criteria required a high likelihood of AD (Braak stage 5 or 6 and CERAD frequent plaques) or intermediate likelihood of AD (Braak stage 3 or 4 and CERAD moderate plaques). Global cognitive function at last visit, summary clinical diagnosis at death, neuritic plaques (CERAD score), and neurofibrillary pathology (Braak stage) are included in **Extended Data Figure 1d–g**, respectively. Clinical and pathologic methods were previously reported.^24,58,59^

As part of the Rush MAP study, blood was drawn at the study home visit and pipetted into two 8.5 mL tiger top serum-separating tubes. Tubes were kept upright for 30 min to 2 h to clot, centrifuged at 1,318 × g for 10 min, aliquoted into 500 µL volumes, and stored at −80 °C. CSF was collected postmortem at autopsy by inserting a needle through the corpus callosum and into the ventricle to extract 10–20 mL. CSF was centrifuged at 1,318 × g for 20 min, aliquoted into 1 mL volumes, and stored at −80 °C. The brain was removed according to standardized protocol and bisected into right and left hemispheres. One hemisphere was fixed and the other hemisphere frozen. The hemisphere to be frozen was slabbed fresh into 1 cm coronal slabs at the time of autopsy. Slabs were placed on pre-cooled (-80 °C) brass plates over dry ice to freeze the tissue rapidly, then placed in resealable bags and stored at −80 °C.

### Mouse cohort design and sample collection

All mouse experiments were conducted in accordance with National Institutes of Health (NIH) guidelines and were approved by the Committee on Animal Care at the Massachusetts Institute of Technology. Mouse colonies were bred and aged in the MIT vivarium with standard light-dark cycles (12 h light/12 h dark), temperature ranges (18–26 °C), and humidity ranges (30–70%) described in the Guide for the Care and Use of Laboratory Animals.^60^ Human *APOE* targeted replacement mice homozygous for *APOE2*, *APOE3*, or *APOE4* were acquired from Taconic Biosciences (strain no. 1547, 1548, and 1549, respectively). In these mouse models, the endogenous mouse locus was replaced with human *APOE* on the C57BL/6NTac background, such that human *APOE* expression remains under control of the endogenous mouse *Apoe* promoter.

The mouse cohort was designed to compare young (3 months) vs. aged (12 months) mice; *APOE2/2, APOE3/3*, vs. *APOE4/4* genotypes; and female vs. male sex. A minimum of *n* = 6 mice were included per group. Blood plasma, CSF, and brain tissue were sequentially collected as follows:

Blood samples were collected by puncturing the submandibular vein. Blood was collected directly into EDTA-coated microvettes (Sarstedt, cat. no. 16.444.100). No more than 10% of the animal’s total blood volume was collected per institutional guidelines. The microvette was gently inverted to mix, stored on ice for no longer than 1 h, then centrifuged at 4,000 × g for 10 min at 4 °C.^61^ Supernatant was collected, leaving the pellet undisturbed, transferred to a clean 1.5 mL microcentrifuge tube, and stored at −80 °C. Plasma samples with abnormal coloration, such as cherry pink indicative of hemolysis, were noted and excluded from analysis.

To collect CSF, mice were anesthetized with ketamine (100 mg/kg) and xylazine (10 mg/kg) administered intraperitoneally. Depth of anesthesia was assessed using the hindlimb pedal withdrawal reflex. Mice remained anesthetized and alive throughout the CSF collection procedure. Mice were secured on a platform with the head flexed forward at a 90° angle. The scalp was disinfected three times with alternating 70% ethanol and betadine, followed by a final 70% ethanol swab, and dissected to access the cisterna magna as follows: a cervical midline incision was made from the sagittal suture to the nape of the neck, and connective tissue and overlying muscles were retracted to expose the atlanto-occipital membrane. Glass capillary needles (Sutter Instrument, cat. no. B100-75-10) were prepared in advance using a micropipette puller (Sutter Instrument, model P-97). Immediately prior to use, capillary needles were opened by trimming the tip with forceps, cleaned with 70% ethanol, and inserted into silicone tubing (1 mm I.D., 1.5 mm O.D.) connected to a 100 µL Gastight Hamilton syringe with a Luer-lock fitting. The capillary needle was flushed with air multiple times then used to gently puncture the atlanto-occipital membrane perpendicularly while maintaining positive pressure during insertion. CSF was collected by slowly withdrawing the syringe plunger, with typical volumes in the range 5–15 µL (average approximately 10 µL). CSF was transferred to a clean 1.5 mL microcentrifuge tube on ice and centrifuged within 1 h, at 4,000 × g for 10 min at 4 °C. Supernatant was transferred to a clean 1.5 mL microcentrifuge tube and stored at −80 °C. Samples were discarded if any blood contamination was observed during sample collection or a red pellet formed after centrifugation.

To collect brain tissue after CSF, mice were further anesthetized by exposure to gaseous isoflurane then transcardially perfused with ice-cold 1× phosphate-buffered saline (PBS). Brains were removed, one hemisphere was snap-frozen on dry ice, and the contralateral hemisphere was immersion-fixed in 4% paraformaldehyde (PFA; diluted from 32% PFA in PBS, Electron Microscopy Sciences, cat. no. 15714-1) for 24 h at 4 °C.

### Lipidomics of MAP human tissue and biofluid samples

For the MAP brain tissue samples, approximately 10 mg of frozen PFC was dissected on a bed of dry ice, transferred to a 1.5 mL microcentrifuge tube, and weighed. All lipidomics sample processing was done in Safe-Lock tubes (Eppendorf, cat. no. 022363204). 200 µL of sterile, cold Dulbecco’s phosphate-buffered saline without magnesium or calcium (DPBS; Gibco, cat. no. 14190144) was added and the sample was homogenized on ice (Cole-Parmer, Motorized Pestle Mixer) for 5 min. An additional 150 µL of sterile, cold DPBS was added and homogenization on ice was repeated for another 5 min. Brain homogenate was then diluted with DPBS to 5 mg/mL frozen-tissue weight per volume, to a total volume of 1 mL. CSF was run neat. Blood serum was 50-fold diluted in sterile, cold water (Sigma-Aldrich, HPLC grade). All tissue samples were visually checked to ensure no clumps or clots prior to re-freezing to −80 °C. One brain and one CSF sample that were visually discolored, indicating likely blood contamination, were excluded.

Mass spectrometry-based lipid analysis was performed by Lipotype GmbH (Dresden, Germany) as previously described.^62^ Lipids were extracted using a chloroform/methanol procedure.^63^ Lipids were extracted from 25–100 µL of homogenized brain tissue (optimized per sample and accounted for in data analysis), 75 µL of neat CSF, or 50 µL of 50-fold diluted blood serum. Samples were spiked with internal lipid standard mixture containing: cardiolipin 14:0/14:0/14:0/14:0 (CL), ceramide 18:1;2/17:0 (Cer), diacylglycerol 17:0/17:0 (DAG), hexosylceramide 18:1;2/12:0 (HexCer), lyso-phosphatidate 17:0 (LPA), lyso-phosphatidylcholine 12:0 (LPC), lyso-phosphatidylethanolamine 17:1 (LPE), lyso-phosphatidylglycerol 17:1 (LPG), lyso-phosphatidylinositol 17:1 (LPI), lyso-phosphatidylserine 17:1 (LPS), phosphatidate 17:0/17:0 (PA), phosphatidylcholine 15:0/18:1 D7 (PC), phosphatidylethanolamine 17:0/17:0 (PE), phosphatidylglycerol 17:0/17:0 (PG), phosphatidylinositol 16:0/16:0 (PI), phosphatidylserine 17:0/17:0 (PS), cholesterol ester 16:0 D7 (CE), sphingomyelin 18:1;2/12:0;0 (SM), triacylglycerol 17:0/17:0/17:0 (TAG) and cholesterol D6 (Chol). After extraction, the organic phase was transferred to an infusion plate and dried in a speed vacuum concentrator. The dry extract was resuspended in 7.5 mM ammonium formate in chloroform/methanol/propanol (1:2:4 vol/vol/vol). All liquid handling steps were performed using Hamilton Robotics STARlet robotic platform with the Anti Droplet Control feature for organic solvents pipetting.

Samples were analyzed by direct infusion on a QExactive mass spectrometer (Thermo Scientific) equipped with a TriVersa NanoMate ion source (Advion Biosciences). Samples were analyzed in both positive and negative ion modes with a resolution of R_m/z=200_=280,000 for MS and R_m/z=200_=17,500 for MS/MS experiments, in a single acquisition. MS/MS was triggered by an inclusion list encompassing corresponding MS mass ranges scanned in 1 Da increments.^64^ Both MS and MS/MS data were combined to monitor CE, Chol, DAG, TAG, and TAG O-ions as ammonium adducts; LPC, LPC O-, PC, and PC O-as formate adducts; and CL, LPS, PA, PE, PE O-, PG, PI, and PS as deprotonated anions. MS only was used to monitor LPA, LPE, LPE O-, LPG, and LPI as deprotonated anions, and Cer, HexCer, and SM as formate adducts.

Data were analyzed with in-house developed lipid identification software based on LipidXplorer.^65,66^ Data post-processing and normalization were performed using an in-house developed data management system. Only lipid identifications with a signal-to-noise ratio >5 and a signal intensity 5-fold higher than in corresponding blank samples were considered for further data analysis. Two PFC and two CSF samples were run as bridge samples across cohorts, however, one PFC sample was excluded due to QC failure.

### Lipidomics of mouse tissue and biofluid samples

Analyses of polar and nonpolar plasma lipids were conducted using an LC-MS system consisting of a Shimadzu Nexera X2 U-HPLC (Shimadzu Corp.) coupled to an Exactive Plus Orbitrap mass spectrometer (Thermo Fisher Scientific). Brain samples were homogenized at a ratio 1 mg tissue per 4 µL water, using a Qiagen TissueLyzer II with 2 × 3 mm tungsten beads at 20 Hz for 4 min. Homogenized brain (10 µL), CSF (10 µL), or plasma (10 µL) samples were extracted with 190 µL of isopropanol containing 1,2-didodecanoyl-sn-glycero-3-phosphocholine (Avanti Polar Lipids) as an internal standard.

After centrifugation, supernatants were injected directly onto a 100 × 2.1 mm, 1.7 µm ACQUITY BEH C8 column (Waters). Volumes injected on the column were brain (2 µL), CSF (10 µL), or plasma (2 µL). The column was eluted isocratically with 80% mobile phase A (95:5:0.1 vol/vol/vol 10 mM ammonium acetate/methanol/formic acid) for 1 min followed by a linear gradient to 80% mobile phase B (99.9:0.1 vol/vol methanol/formic acid) over 2 min, a linear gradient to 100% mobile phase B over 7 min, then 3 min at 100% mobile phase B. MS analyses were carried out using electrospray ionization in the positive ion mode using full scan analysis over 200–1100 m/z at 70,000 resolution and 3 Hz data acquisition rate. Other MS settings were: sheath gas 50, in-source CID 5 eV, sweep gas 5, spray voltage 3 kV, capillary temperature 300 °C, S-lens RF 60, heater temperature 300 °C, microscans 1, automatic gain control target 1 × 10^6^, and maximum ion time 100 ms.

Raw data were processed using TraceFinder software (Thermo Fisher Scientific) for targeted peak integration and manual review of a subset of identified lipids and using Progenesis QI (Nonlinear Dynamics) for peak detection and integration of both lipids of known identity and unknowns. Mixtures of reference standards and reference samples were included in each analysis queue to confirm IDs in every dataset. For cross-compartment comparisons, feature alignment was performed using a custom software tool to detect landmark features and non-parametric retention time scaling to accurately match unknowns between datasets. Each sample was median-normalized. Lipid identities were determined based on comparison to reference plasma extracts and are denoted by total number of carbons in the lipid acyl chain(s) and total number of double bonds in the lipid acyl chain(s). All tissues were nearest-neighbor normalized.

### Lipidomics data analysis

The lipidomic data was analyzed after having undergone data standardization, feature alignment, quality control, and metabolite annotation, as described above for the MAP and mouse cohorts. All subsequent analysis was completed in Python using standard packages including SciPy, NumPy, statsmodels, pandas, and Scikit-learn. Three human brain samples were excluded from further analysis due to injection normalization errors. Brain lipid abundances were normalized by the ratio of tissue concentration (5 µg/µL) to sample volume. Unannotated mouse lipids were not included except in cross-fluid PCA and correlation analysis. Lipids detected in three or fewer samples per set were also excluded. For the remaining lipids, missing values were imputed as half the minimum detected value of each feature within each sample set and tissue. This approach limits the number of lipids excluded due to lack of detection in a particular diagnostic group.^67^ After processing each MAP sample set separately, the two sets were combined for downstream analyses, retaining only lipids that met the detection threshold in both sets. For pairwise analyses between groups (e.g., volcano plots), lipids detected in fewer than three samples in both groups were additionally excluded. For cross-compartment PCA, lipids were z-scored across all samples and only lipids that were present in all sample data after post-processing were included. For heatmaps, the single-compartment PCA, and the cholesterol comparison plots, scores were normalized within each tissue.

Fractional contribution of each lipid class to the total lipid content was calculated as the median of the sum of lipid abundances within each class across individuals, normalized to the sum of class medians. To quantify relative composition shifts, CLR_2_ was calculated as the log_2_ of a feature in a sample divided by the geometric mean of all features in the sample. For human brain and CSF, a linear model was used to control for sample set effects when calculating fold-change and ΔCLR_2_. *P* values on coefficients for lipid abundance fold-change, ΔCLR_2_, and fold-change of summed class totals were calculated using a two-sided t-test for independent samples with homoscedastic error (equivalent to an equal-variance t-tests when no sample set term is included). Benjamini-Hochberg correction was additionally applied to determine false-discovery-rate (FDR)-adjusted *P* value threshold of 0.05, as indicated in the volcano plots. log_2_(fold-change) is computed as the difference in mean log_2_ lipid abundances between comparison groups. To evaluate population similarity, MAP donor groups were clustered using a hierarchical dendrogram based on normalized lipid abundance profiles with Euclidean distance and median linkage. Ordinary least-squares linear regressions quantified the relationship between variables, including enrichment vs. number of lipid double bonds and CSF cholesterol sample z-scores vs. that in the brain, with *P* values of the coefficients determined using a two-sided t-test without false-discovery-rate corrections. In CSF, log_2_-transformed lipid class totals per individual were regressed against group properties controlling for sample set as:

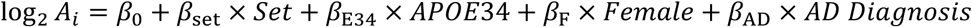

To assess the correlation of cognitive level with donor parameters, we z-score normalized and separately regressed global cognition at last visit against the normalized amyloid content in the brain and lipid species log_2_ abundances controlling for sample set. Brain amyloid content represents the percent area of the cortex occupied by amyloid measured by immunohistochemistry and averaged across eight brain regions.

Lipids identified in the mouse lipidomics dataset predominantly did not resolve individual acyl chains, whereas the human lipidomics dataset did. To compare lipidomic findings between species, lipids with designated acyl groups were reduced to the notation of class + total carbons:double bonds, with abundances summed for lipids mapping to the same resulting identifier; isomers were merged, and hydroxyl or oxidation marks were dropped. Correlations between lipid changes in human and mouse data were quantified using Spearman correlation coefficients with corresponding asymptotic *P* values.

### rLP synthesis and purification

Lipids (Avanti Research) were supplied in or solubilized to 10 mg/mL (phospholipids) or 100 mg/mL (cholesterol) in chloroform, and stored in glass vials with PTFE-lined caps at -20 °C. Lipids included: 1-palmitoyl-2-oleoyl-glycero-3-phosphocholine, or 16:0-18:1 PC (POPC; Avanti Research, cat. no. 850457C); 1-palmitoyl-2-oleoyl-sn-glycero-3-phosphoethanolamine, or 16:0-18:1 PE (POPE; Avanti Research, cat. no. 850757C); 1-stearoyl-2-oleoyl-sn-glycero-3-phosphoethanolamine, or 18:0-18:1 PE (SOPE; Avanti Research, cat. no. 850758P); and cholesterol (Avanti Research, cat. no. 700100P). Sodium cholate (Sigma-Aldrich, cat. no. C6445) was solubilized to 200 mM in 1× PBS and stored at room temperature in the dark. ApoE (PeproTech, cat. no. 350-02 for ApoE3 and 350-04 for ApoE4) was reconstituted to 1 mg/mL in PBS and stored at -20 °C.

rLPs were synthesized via thin-film hydration and cholate-mediated ApoE incorporation using an initial molar ratio of 70:17:8:5:1 for POPC:SOPE:POPE:cholesterol:ApoE, at a scale of 800 µg total lipid and ApoE per vial. Specifically, lipids in chloroform were combined using Hamilton Gastight syringes into 2 mL glass vials. Where indicated, a lipophilic dye was also incorporated at this stage, Vybrant DiI (Invitrogen, cat. no. V22885; 2% by mass relative to total phospholipid content). To facilitate scalable preparation, the lipid mixture was dried under vacuum using a Centrifuge Concentrator (Eppendorf, Vacufuge plus) in the vacuum, high vapor mode at 30 °C for 2 h, forming a thin lipid film on the glass vial. Sodium cholate was diluted in PBS to 50 mM and added at a molar ratio of 3:1 to the total lipid content to rehydrate the lipid film. The mixture was bath-sonicated in 10-s intervals until the lipid film was fully dissolved, approximately 5–10 min. After the solution appeared homogeneous and clear, ApoE was added and gently mixed by pipetting. More sodium cholate was added until its final concentration was 20 mM (above the critical micelle concentration). The solution was transferred to a 1.5 mL microcentrifuge tube and mixed on a tube rotator at a temperature above the maximum lipid transition temperature (T > T_m_; 37 °C) for at least 12 h. Note all rLP synthesis, purification, and downstream handling were performed in Protein LoBind tubes (Eppendorf, cat. no. 0030108116). Samples were then dialyzed for 24 h at the same incubation temperature (T > T_m_; 37 °C), with a buffer change at approximately the halfway point, using either 10 kDa MWCO Slide-A-Lyzer 0.1–0.5 mL cartridges (Thermo Fisher Scientific, cat. no. 66384) against 1.8 L of PBS with gentle stirring, or the replacement 10 kDa MWCO Slide-A-Lyzer MINI Dialysis Devices (Thermo Fisher Scientific, cat. no. 88401) against 14 mL of PBS with gentle orbital shaking.

Samples were centrifuged at 10,000 × g for 15 min at 4 °C to pellet any precipitants, then supernatant was concentrated by 30 kDa MWCO Amicon® ultrafiltration (Millipore Sigma, cat. no. UFC503096; 15 min at 14,000 × g followed by 2 min at 1,000 × g reverse spin). rLP product was isolated via size-exclusion chromatography (SEC) using Superdex™ 200 Prep Grade resin (Cytiva, cat. no. 17104301) in 0.5–0.7 × 15 cm columns (Bio-Rad, Econo-Column® Chromatography Columns, cat. no. 7370517 or 7370717) via gravity-driven flow. Resin was loaded following manufacturer’s instructions. Columns were flushed overnight with PBS prior to use and the sample was loaded. The first fraction was collected after 30 min, followed by eight subsequent fractions at 15-min intervals. Fractions containing rLPs were identified based on absorbance at 280 nm (protein) and 550 nm where applicable (lipophilic dye), and hydrodynamic diameter measured by DLS (see characterization methods below). Fractions were stored in the dark at 4 °C and used within 2 weeks, and prior to use, were concentrated by 30 kDa MWCO ultrafiltration, diluted to the required final concentration, and sterile filtered using 0.22 µm Millex® PVDF syringe filter (Millipore, cat. no. SLGV004SL).

### Absorbance characterization of rLPs

To determine SEC fractions containing product, absorbance of ApoE and Vybrant DiI was measured at 280 nm and 550 nm, respectively, using a NanoDrop spectrophotometer (Thermo Fisher Scientific) with baseline correction at 340 nm. The final concentration of purified, concentrated rLP product was determined by similarly measuring absorbance and converting to concentration using the extinction coefficients of 44,460 M^−1^ cm^−1^ for ApoE and 148,000 M^−1^ cm^−1^ for Vybrant DiI. All subsequent rLP concentrations are reported based on ApoE protein concentration.

### Hydrodynamic size and zeta potential characterization of rLPs

Fractions with maximal ApoE and DiI present were analyzed via dynamic light scattering (DLS) to confirm the hydrodynamic diameter was in the anticipated size range. DLS and zeta potential measurements were completed on the Malvern Panalytical Zetasizer Nano with the diluent set as water, the material as liposomes, and 60 s equilibration time prior to the run. For DLS, 400 µL volume of undiluted rLP or 0.1 mg/mL ApoE was measured in a disposable cuvette (BrandTech Scientific, cat. no. 759150). Zeta potential was measured as a proxy for surface charge and particle stability by diluting samples to 2.5 µg/mL protein concentration in water and loading 1 mL volume in a folded capillary zeta cell (Malvern Panalytical, cat. no. DTS1070).

### TEM characterization of rLPs

For negative-stain room-temperature transmission electron microscopy (RT TEM), 10 µL of sample at 0.1 mg/mL in PBS was applied to a 200-mesh copper grid coated with a continuous carbon film. After 60 s, excess solution was removed by gently touching the grid edge with a Kimwipe. Next, 10 µL of 2% aqueous uranyl acetate (negative stain) was applied to the grid. After 30 s, excess stain was similarly removed, and the grid was air-dried at room temperature. The prepared grid was mounted on a JEOL single-tilt holder and inserted into the TEM column.

For cryogenic transmission electron microscopy (cryo-TEM), 3 µL of the sample at 1.5 mg/mL in PBS was applied to a lacey copper grid coated with a continuous carbon film. Excess sample was gently blotted using the Gatan Cryo Plunge III, followed by rapid plunging into liquid ethane to vitrify the sample. The prepared grid was mounted on a Gatan 626 single-tilt cryo-holder and inserted into the TEM column. Both the specimen and holder tip were maintained at cryogenic temperatures with liquid nitrogen to ensure preservation throughout the transfer and imaging process.

Imaging for both RT and cryo-TEM was performed on a JEOL 2100 FEG microscope using a minimum dose method to minimize electron beam-induced damage. The microscope was operated at 200 kV, with magnifications ranging from 10,000× to 60,000× to assess particle size and distribution. All images were captured using a Gatan 2k × 2k UltraScan CCD camera.

Particle dimensions were measured manually in Fiji (ImageJ2, v2.14.0/1.54g) using the straight-line tool on TEM images calibrated to the embedded scale bar. Particles were classified by orientation: face-on particles (circular) were measured for diameter, and edge-on particles (oblong) for both long-axis diameter and short-axis, apparent bilayer thickness. Face-on and edge-on particle diameters matched. Ten particles of each orientation were measured in each of four fields of view, from a single rLP preparation.

### Lipid and amyloid-β binding assays in solution

Lipid binding capacity of rLPs was determined using nitrobenzoxadiazole (NBD) fluorophore-labeled lipid analogues. NBD is weakly fluorescent in polar environments (free in aqueous solution) and becomes more fluorescent in nonpolar environments (upon rLP binding). Three NBD-lipids were tested: NBD palmitic acid (NBD-C16:0; Avanti Research, cat. no. 810105), C12 NBD Galactosylceramide (d18:1/12:0) (NBD-GalCer; Cayman Chemical, cat. no. 22851), and 25-NBD-cholesterol (Avanti Research, cat. no. 810250). NBD-lipids were reconstituted in DMSO to 0.2 mg/mL, aliquoted, and stored at -20 °C. rLPs (synthesized without DiI lipophilic dye) were diluted to a stock concentration of 20 µg/mL in PBS. rLPs and NBD-lipids were mixed in microcentrifuge tubes to a final concentration of 5 or 10 µg/mL rLPs and varying amounts of NBD-lipids ranging from 0 to 25 µM final concentrations, diluted in PBS. A separate set of NBD-lipids alone (no rLPs) was prepared at the same concentrations to calculate background fluorescence in aqueous solution. Samples were vortexed for 5 s, briefly centrifuged, and incubated for 20 min at room temperature. Volumes of 50 µL were transferred into a flat clear bottom black-sided 384-well plate (Corning, cat. no. 3764). Fluorescence spectra were measured using a plate reader (Tecan, Infinite® M Plex) with excitation at 473 nm and emission scanned from 510 to 650 nm in 1 nm increments with 20 µs integration time.

Amyloid-β peptide binding capacity of rLPs was determined as described above with the following modifications: Aβ42-HiLyte Fluor 647 (AnaSpec, cat. no. AS-64161) was reconstituted in 1% (v/v) ammonium hydroxide to 1 mg/mL, immediately diluted to 20 µM in PBS, aliquoted, and stored at -20 °C; Aβ42-HiLyte Fluor 647 final concentrations ranged from 0 to 0.5 µM; after incubation with or without rLPs, unbound Aβ42 was removed by 30 kDa MWCO Amicon® ultrafiltration (Millipore Sigma, cat. no. UFC503096; 15 min at 14,000 × g) and the retentate was measured. Fluorescence spectra were measured with excitation at 600 nm and emission scanned from 650 to 820 nm in 1 nm increments with 20 µs integration time.

NBD or HiLyte fluorescence change (relative to the control without rLPs) was calculated at the approximate maxima emission wavelengths of 544 or 674 nm, respectively and fit using a site-binding isotherm:

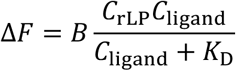

where *C*_rLP_ and *C*_ligand_ are the molar concentrations of the added rLPs and ligands, respectively, *K*_D_ is the site dissociation constant, and *B* is a constant that is the product of the number of binding sites per rLP and the change in fluorescence of the free to bound ligand dye. The fitting constants were determined by minimizing the sum of squared residuals fit of the fluorescence change, with the standard errors from the covariance matrix using the curve-fit function in scipy across all concentrations of rLP and ligand.

### Cholesterol efflux assay in human iPSC-derived astrocytes

Human induced pluripotent stem cell (iPSC) lines were generated by the Picower Institute for Learning and Memory iPS Core Facility. *APOE4* iPSCs used in this study originated from line AG09173 (homozygous *APOE3* parental, female donor, age 75 at biopsy) that was CRISPR/Cas9-edited to homozygous *APOE4* as previously described.^16^ iPSCs were maintained in mTeSR™1 medium (STEMCELL Technologies, cat. no. 85850) on plates coated with 2% (v/v) Matrigel® hESC-Qualified Matrix (Corning, cat. no. 354277) diluted in the same medium and passaged at approximately 60% confluence using ACCUTASE™ (STEMCELL Technologies, cat. no. 07920).

Astrocytes were differentiated from iPSCs as described previously,^68^ plated onto 96-well glass-bottom plates (CellVis, cat. no. P96-0-N) using the same Matrigel coating conditions described above at a density of 1.5 × 10^4^ cells per well, and cultured in astrocyte medium (ScienCell Research Laboratories, cat. no. 1801) with medium changed every-other day for four days or until confluence. Experiments were performed in serum-free, phenol-red-free astrocyte medium (ScienCell Research Laboratories, cat. no. 1801-prf without FBS supplement). To visualize intracellular cholesterol, cells were incubated with 200 µL of medium with 1 µg/mL BODIPY 480/508-Cholesterol (Cayman Chemical, cat. no. 24618) at 37 °C, 5% CO_2_ for 1 h. Cells were washed with medium once, then incubated with 100 µL of medium with either PBS (control) or 10 µg/mL sterile-filtered rLPs, at 37 °C, 5% CO_2_ for 24 h. To investigate cholesterol-efflux pathways, BODIPY-cholesterol-loaded astrocytes were co-incubated with PBS (control) or rLPs (10 µg/mL) together with probucol (20 µM; Sigma-Aldrich, cat. no. PHR3112), GSK2033 (10 µM; MedChemExpress, cat. no. HY-108688), or DMSO (vehicle) for 24 h (*n* = 3 wells per condition).

After 24 h, 80 µL of spent medium was collected and transferred into a flat clear bottom black-sided 384-well plate (Corning, cat. no. 3764) to measure levels of BODIPY-cholesterol and rLPs in the medium with a fluorescence plate reader (Tecan Infinite M Plex; excitation/emission wavelengths of 480/515 nm for BODIPY-cholesterol and 550/610 nm for DiI in rLPs). Remaining medium was aspirated and cells were washed once with medium prior to imaging. Live cells were imaged (Agilent BioTek Cytation C10 confocal imaging reader) with the imaging chamber maintained at 37 °C, 5% CO_2_ incubation conditions. Imaging was performed with a 20× objective (Olympus, Plan Fluorite), 40-µm Nipkow disk, with excitation/emission wavelengths of 472/520 nm for BODIPY-cholesterol and 556/600 nm for DiI in rLPs, across a z-stack containing the full height of the cells. Experiments were conducted with *n* = 3 wells per condition, and *n* = 4 regions of interest (ROIs) per well.

BODIPY-cholesterol was analyzed using a custom automated script in Fiji (ImageJ2, v2.14.0/1.54g). Briefly, optimal threshold values were manually determined for each batch based on a subset of control and treated images. Using the automated script, each image was opened, thresholded to these settings, and converted into a binary mask. Lipid droplet counts were calculated with the Analyze Particles function, applying size criteria of 0.1–12.5 µm^2^ (equivalent to diameter bounds of ∼360 nm – 4 µm).^69–71^

### shRNA-mediated knockdown of *ABCA1* and *ABCG1*

Lentiviral vectors expressing U6 promoter–driven shRNAs targeting human *ABCA1* (VectorBuilder, VB260304-1354rjc; 10.02 × 10^8^ TU/mL) or *ABCG1* (VectorBuilder, VB260304-1363uze; 9.39 × 10^8^ TU/mL), each carrying a neomycin resistance cassette (pLV[shRNA]-Neo-U6), were packaged by VectorBuilder. A non-targeting scrambled shRNA in the same backbone (VectorBuilder, VB010000-9536ysn; 7.34 × 10^8^ TU/mL) served as the control.

Human *APOE4* iPSC-derived astrocytes generated as described in the previous section were plated on day 0 at 5 × 10^4^ cells per well in Matrigel-coated 24-well plates, with medium changed every-other day. On day 3, astrocytes were transduced at a multiplicity of infection of 10, calculated from the day-0 cell number, astrocyte medium (ScienCell Research Laboratories, cat. no. 1801-prf). Medium was fully replaced on day 5, and cells were cultured for an additional 2 days. On day 8, knockdown of *ABCA1* and *ABCG1* was validated by RT-qPCR. Total RNA was isolated using the High Pure RNA Isolation Kit (Roche Diagnostics) following the manufacturer’s instructions, and RNA concentration and purity were assessed by NanoDrop spectrophotometry (A260/280 and A260/230 ratios; acceptable range 1.8–2.2). cDNA was synthesized from 200 ng RNA in 20 µL reactions using iScript Reverse Transcription Supermix (Bio-Rad, cat. no. 1708841). The entire cDNA reaction was diluted 1:5 in nuclease-free water (2 ng/µL RNA equivalents), and 2 µL (4 ng RNA-equivalent cDNA) was used per quantitative PCR reaction. Reactions were run in technical duplicate in 20 µL, containing 10 µL of 2× SsoAdvanced Universal Probes Supermix (Bio-Rad, cat. no. 1725281), 1 µL of 20× TaqMan Gene Expression Assay and 2 µL of diluted cDNA, using assays for *ABCA1* (Hs01059101_m1), *ABCG1* (Hs00245154_m1), and *ACTB* (Hs01060665_g1) as the reference. Reactions were run on a Bio-Rad CFX96, and relative expression was calculated using the 2⁻ΔΔCt method, with knockdown expressed relative to scrambled-shRNA controls.

On day 7, cells were loaded with 1 µg/mL BODIPY 480/508-cholesterol at 37 °C, 5% CO_2_ for 1 h, followed by the cholesterol-efflux assay described above. Cells were treated with PBS (control), free ApoE3, or rLPs (2.5 µg protein per well) in 250 µL of medium for 24 h, after which medium in duplicate was collected and fluorescence was quantified.

### Bulk RNA sequencing of human iPSC-derived astrocytes with rLPs

Human iPSC-derived *APOE4* astrocytes were differentiated and cultured as described above, then treated with either serum-free medium alone (control; *n* = 5) or 10 µg/mL rLPs diluted in the same medium (*n* = 5) at a volume of 1 mL per well in 6-well plates for 24 h. Following treatment, cells were washed once with DPBS, dissociated with ACCUTASE™ for 5–10 min, and centrifuged at 300 × g for 5 min. Supernatant was aspirated, and the cell pellet was resuspended in 300 µL TRIzol (Invitrogen, cat. no. 15596018) and incubated at room temperature for 5 min. RNA was extracted using the Direct-zol RNA Miniprep Kit (Zymo Research, cat. no. R-2062) following manufacturer’s instructions.

RNA quality was assessed using a Fragment Analyzer (Agilent Technologies). RNA-seq libraries were prepared from 100 ng total RNA using the NEBNext Poly(A) mRNA Magnetic Isolation Module (New England Biolabs, cat. no. E7490) and NEBNext Ultra II RNA Library Prep Kit for Illumina (New England Biolabs, cat. no. E7770), following the manufacturer’s instructions with 14 cycles of PCR and Singular-compatible combinatorial indexes. Final libraries were sized using the Fragment Analyzer, quantified by qPCR, normalized, and sequenced on a Singular G4 using a paired-end configuration of 2 × 50 bp with dual 8-bp index reads.

Sequencing quality was assessed using FastQC (v0.12.0). Reads were aligned to the human reference genome (hg19) using STAR (v2.7.11), and gene-level counts were generated using featureCounts from the Subread package (v2.1.1).

Genes with ≥10 counts in at least two samples were retained. Counts were TMM-normalized using edgeR (v3.36.0) and analyzed with limma-voom (limma v3.50.3) using a treatment-group model and empirical Bayes moderated t-tests. P-values were adjusted using the Benjamini-Hochberg method. Genes with adjusted *P* < 0.05 and |log₂ fold-change| ≥ 0.5 were classified as differentially expressed in volcano plots.

Gene set enrichment analysis was performed using fgsea (v1.20.0). Pathways with adjusted *P* < 0.05 were considered significantly enriched. Selected cholesterol-related and extracellular matrix organization leading-edge genes were highlighted in separate volcano plots.

### Intracisternal injection of rLPs in mice

General mouse study information and procedures are described in the section “Mouse cohort design and sample collection.” 5xFAD transgenic mice were acquired from the Mutant Mouse Resource and Research Centers (MMRRC; stock no. 034848-JAX; distributed by The Jackson Laboratory) and human *APOE4* targeted-replacement mice were acquired from Taconic Biosciences (strain no. 1549). 5xFAD and *APOE4* mice were crossed for at least eight generations.

*APOE4*;5xFAD mice (7–8 months old for behavior experiments or 12–14 months old for IHC and -omics characterization) were weighed and placed into an isoflurane induction chamber at 2.5– 3% isoflurane until hindlimb pedal withdrawal reflexes ceased. Mice were subsequently transferred to a nose cone to sustain isoflurane anesthesia at 1.5–2% and fur was shaved from the occipital crest to the nape. Meloxicam (Meloxivet, 5 mg/mL solution) and buprenorphine sustained-release were subcutaneously administered at 5 mg/kg and 1 mg/kg, respectively. Mice were next placed into a stereotaxic frame over a heating pad, maintaining isoflurane delivery through a mouth bar at 1.5–2%. Ophthalmic ointment was applied to eyes, and 2.5% lidocaine/2.5% prilocaine cream was applied to ear bar insertion points before fixing the head to the stereotaxic frame. Shaved skin was disinfected (three times alternating 70% ethanol then betadine, followed by a final 70% ethanol swab), an incision was made from the occipital crest to the nape, and skin and connective tissue were separated to expose muscles using fine-pointed forceps. Using scissors and fine-pointed forceps, vertical incisions of muscle tissue were made, and tissue was pulled aside to allow puncture into the cisterna magna. A clean Nanofil syringe (World Precision Instruments, cat. no. NC0578818) with an attached Nanofil needle (World Precision Instruments, cat. no. NF34BV) was filled with 10 µL of either sterile saline (control) or 0.5 mg/mL sterile-filtered rLPs diluted in sterile saline, and secured onto a microinjection syringe pump (World Precision Instruments, model UMP3). Using the stereotaxic frame, the syringe was positioned and lowered until the needle punctured the atlanto-occipital membrane. The needle was lowered as deeply as possible into the cisterna magna without contacting brain tissue. Saline or rLPs were injected into the cisterna magna using the syringe pump at a rate of 1 µL/min, and the insertion point was monitored to ensure minimal backflow during injection. 10 min after cessation of injection, the needle was retracted slowly at 0.1 mm/s and the incision was sutured. Mice were placed into their home cage and monitored during recovery until alert and ambulatory, ensuring the absence of motor defects.

At the experimental endpoint (24 h for acute studies or 30 days for behavior experiments), mice were anesthetized with gaseous isoflurane (USP) and transcardially perfused using a perfusion pump (MSE Supplies, cat. no. LS0216) and 25G × ¾-inch winged infusion set needles (Surflo, 0.5 × 19 mm). For snRNA-seq or lipidomic analyses, mice were perfused with ice-cold PBS, brains were removed, and the hippocampus and prefrontal cortex from one hemisphere were dissected on ice, snap-frozen on dry ice, and stored at −80 °C. The contralateral hemisphere was immersion-fixed in 5 mL of 4% paraformaldehyde (PFA; diluted from 32% PFA in PBS, Electron Microscopy Sciences, cat. no. 15714-1) in a 15-mL conical tube overnight at 4 °C for immunohistochemistry (IHC). For whole-brain IHC, mice were perfused with chilled PBS for 7 min followed by 4% PFA for an additional 7 min. Brains were harvested and similarly immersion-fixed in 4% PFA overnight at 4 °C.

### Mouse immunohistochemistry

Following fixation as described above, brains were transferred into 12 mL of 30% sucrose (Sigma-Aldrich, cat. no. S7903-KG) and incubated for 3 days at 4 °C, or until the tissue had fully sunk. Brains were then sectioned coronally into 40-µm slices using a vibrating blade microtome (Leica Biosystems VT1000S) and stored in PBS at 4 °C. All subsequent incubation steps were performed on an orbital shaker at 50 rpm. Sections were washed with PBS for 10 min at room temperature (RT), then incubated in a blocking and permeabilization solution (BPS) of 5% (w/v) BSA and 0.3% (v/v) Triton X-100 in PBS for 1 h at RT. Sections were then placed in primary antibodies (β-Amyloid D54D2 Rabbit Monoclonal Antibody, Cell Signaling Technology, cat. no. 8243, 1:500; GFAP 2.2B10 Rat Monoclonal Antibody, Invitrogen, Thermo Fisher Scientific, cat. no. 13-0300, 1:750; Iba1 Gp311H9 Guinea Pig Recombinant Monoclonal Antibody, Synaptic Systems, cat. no. 234 308, 1:500) diluted in BPS for 24 h at 4 °C. After three 10-min washes in PBS, sections were incubated in secondary antibodies in BPS for 2 h at RT. Immunostained sections were then washed with PBS for 10 min. For lipid droplet staining, sections were incubated with BODIPY 493/503 (Invitrogen, cat. no. D3922) at a concentration of 1 µg/mL in PBS for 1 h in 24-well mesh-bottom wells to facilitate simultaneous removal of sections, followed by a 10-min PBS wash. Sections were then stained with Hoechst (Invitrogen, cat. no. H3570) at a dilution of 1:10,000 in PBS for 15 min in 24-well mesh-bottom wells, followed by a final 10-min PBS wash. Tissue mounting was performed in PBS on SuperFrost slides (Fisher Scientific, cat. no. 22-037-246) and coverslips were added using Fluoromount-G mounting medium (Southern Biotech, cat. no. 0100-01).

For VGLUT1 and VGAT staining, two sections from each mouse were placed into a 24-well plate (VWR, cat. no. 734-2779), containing 500 µL of PBS (Cytiva, cat. no. SH30256.01) and washed three times with PBS, with 5-minute incubations on a shaker. For blocking, samples were incubated in 300 µL of blocking solution containing MAXblock Blocking (Active Motif, cat. no. 15252) with 0.3% Triton X-100 (Sigma-Aldrich, cat. no. X100-100mL) for 2 hours at room temperature. Primary antibodies, including anti-VGLUT1 (Synaptic Systems, cat. no. 135-302, 1:500) and anti-VGAT (Cl Gp117G4; Synaptic Systems, cat. no. 131-308, 1:500), were diluted in staining solution containing MAXbind Staining (Active Motif, cat. no. 15253) and 0.3% Triton X-100. The antibody solution (300 µL) was added to each well, and samples were incubated overnight at 4 °C on a continuous shaker. Following primary antibody incubation, sections were washed three times with PBS, with 5-minute incubations on a shaker. Secondary antibodies, including Alexa Fluor 555-conjugated anti-rabbit IgG (Invitrogen, cat. no. A-31572, 1:500) and Alexa Fluor 647-conjugated anti-guinea pig IgG (Invitrogen, cat. no. A-21450, 1:500), along with DAPI (Thermo Fisher Scientific, cat. no. 62248, 1:5000), were prepared in staining solution and incubated with the sections (300 µL) for 2 hours, protected from light. Sections were subsequently washed three times with PBS, with 5-minute incubations on a shaker. Finally, stained sections were mounted onto glass slides (25 × 75 × 1.0 mm; Fisherbrand, cat. no. 1255015) and air-dried in the dark for approximately 10 minutes. Using mounting medium (Vector Laboratories, cat. no. H-1000-10), slides were coverslipped with 24 × 40 mm coverslips (Avantor, cat. no. 48393-060).

### Mouse brain imaging and quantification

Imaging was performed on a Zeiss LSM900 confocal microscope with ZEN Blue software (Carl Zeiss Microscopy, v3.9). Uniform acquisition settings were applied across each image set. Images were acquired at 10× (Zeiss, Objective Plan-Apochromat 10×/0.3 M27 thread) and 20× (Zeiss, Objective Plan-Apochromat 20×/0.8), as noted in figure captions. Laser lines included 405, 488, 561, and 640 nm, kept on separate imaging tracks with emission windows set via variable secondary dichroics to avoid spectral overlap (410–470 nm, 492–550 nm, 564–625 nm, and 650– 700 nm, respectively). Laser-blocking filters were also applied on all lines and a shortpass 470 filter on the 405 nm excitation line. Pinhole diameter was set to one Airy unit (1 AU) for each laser line. Laser intensity and detector gain were optimized within each dataset to avoid saturating the detector. Total z-stack height was set to capture the full tissue thickness and optimal z-stack step size was determined by half the z-resolution according to the Nyquist criterion. Tiling regions were set to capture the full region of interest: 3 × 2 for hippocampus (10× objective; 3580 µm × 2430 µm; 0.8014 pixels per µm), 2 × 2 for prefrontal cortex (10× objective; 2430 µm × 2430 µm; 0.8014 pixels per µm), and 1 × 1 for subiculum (20× objective; 638.9 µm × 638.9 µm; 1.6028 pixels per µm).

Analysis was completed using Fiji (ImageJ2, v2.14.0/1.54g) and Imaris (Oxford Instruments, v10.2) software. Researchers analyzing imaging data were blinded to the experimental conditions. Lipid droplets in the subiculum were quantified from BODIPY 493/503-stained z-stacks using a custom Fiji script. The subiculum region of interest (ROI) was manually defined, and batch-specific intensity thresholds were applied. 3D particle analysis identified lipid droplets within a volume range of 0.0335–33.51 µm^3^, and droplet counts and integrated BODIPY fluorescence were extracted. Aβ plaques were quantified in Imaris using machine-learning 3D segmentation of D54D2-positive signal within manually defined hippocampal regions of interest (ROIs) or uniformly cropped square regions in the PFC. For each imaging batch and region, a machine-learning classifier was trained iteratively using five randomly selected images. A seed-point diameter of 10 µm and a minimum plaque volume of 5 µm^3^ were applied for amyloid segmentation. The final classifier was then applied to all images from the corresponding batch and region. Plaque count (number of disconnected components), segmented amyloid volume, and ROI dimensions were exported for analysis. GFAP and IBA1 immunoreactivity was quantified as the percentage of the region-of-interest area occupied by marker-positive pixels using intensity threshold–based pixel classification in QuPath (v0.7.0). Classifier settings included full-resolution analysis and Gaussian prefiltering with a smoothing sigma of 0. Sections were analyzed together, with same classifier applied. Pixels with intensities above the corresponding threshold were classified as GFAP-or IBA1-positive. For each marker, the classified positive area was divided by the total region-of-interest area and expressed as a percentage. Measurements from two brain sections were averaged per mouse before statistical analysis.

VGLUT1 and VGAT immunoreactivity was quantified as the percentage of the region-of-interest area occupied by marker-positive pixels using intensity threshold–based pixel classification in QuPath (v0.7.0). Classifier settings included full-resolution analysis and Gaussian prefiltering with a smoothing sigma of 0.3. Sections were analyzed in two batches, with marker-specific intensity thresholds defined separately for each batch and applied uniformly to all images within that batch. Pixels with intensities above the corresponding threshold were classified as VGLUT1-positive or VGAT-positive. For each marker, the classified positive area was divided by the total region-of-interest area and expressed as a percentage.

### Mouse brain single-nucleus RNA-sequencing (snRNA-seq)

Mice were euthanized and tissue harvested as described above in the section “Intracisternal injection of rLPs in mice”. Nuclei were isolated from frozen hippocampal brain tissue following a previously described protocol,^72^ with some modifications. Briefly, tissue was homogenized in 700 µL homogenization buffer, then the lysate after filtration through a 40 µm cell strainer (Falcon, cat. no. 352340) and addition of 450 μL working solution was loaded as a 25% iodixanol solution on top of a 30%–40% iodixanol density gradient (750 μL 30% Iodixanol solution, 300 μL 40% iodixanol solution). Nuclei were separated by centrifugation on a fixed rotor, tabletop centrifuge (10000 × g for 5 min at 4 °C). The nuclei pellet was collected and transferred to a new 1.5 mL DNA LoBind tube (Eppendorf, cat. no. 0030108.051), washed twice with 1 mL ice-cold PBS/1% BSA buffer, centrifuged ( 300 × g for 3 min at 4 °C), and resuspended in 100 μL PBS/1% BSA buffer. Droplet-based single-nucleus RNA-seq libraries were generated using the Chromium GEM-X Single Cell 3′ Kit v4 (10x Genomics, 16 reactions, cat. no. PN-1000691) on a Chromium X instrument (10x Genomics). Libraries were sequenced through the MIT BioMicro Center on an Illumina NovaSeq 6000 using two S1 flow cells with 28 + 90 bp paired-end reads and 10 + 10 bp dual indexing. Sequencing reads from both flow cells were combined for each sample and aligned to the mouse reference genome GRCm39 and quantified using Cell Ranger (10x Genomics, v9.0.1). Doublets were identified per sample using scDblFinder^73^ and removed. Low-quality nuclei were excluded using the following thresholds: 200–6,000 detected genes per nucleus and <5% mitochondrial reads; genes detected in fewer than 10 cells were removed. Data were normalized with SCTransform (v2, glmGamPoi),^74,75^ regressing out detected gene number, mitochondrial percentage, age, sex, and parental genotype origin. Nuclei were clustered and annotated into major brain cell populations, with vascular cells subclustered separately to resolve endothelial cells, pericytes, vascular smooth muscle cells, and fibroblast-like cells based on canonical marker expression.

Differential expression between treated and control animals was assessed within each annotated cell type at both the single-cell and pseudobulk level. Single-cell testing used Wilcoxon rank-sum tests, with genes classified as up- or downregulated at adjusted *P* < 0.05 and |log_2_ fold-change| > 0.25. For pseudobulk analysis, raw counts were aggregated per sample and cell type. Cell types with ≥4 pseudobulk samples across both groups were retained; genes were filtered with filterByExpr (edgeR),^76^ libraries normalized by TMM, and counts transformed with voom (limma).^77^ Differential expression was assessed using linear models with empirical Bayes moderation, including sex and parental inheritance of 5xFAD transgene as covariates. Functional enrichment was assessed by two complementary approaches. Over-representation analysis of GO Biological Process terms was performed on cell-type-specific differentially expressed gene sets using clusterProfiler^78^ with org.Mm.eg.db, Benjamini-Hochberg correction, and an adjusted *P* value cutoff of 0.05, run separately for up- and downregulated genes; redundant terms were collapsed with simplify (similarity cutoff = 0.7). Gene set enrichment analysis was performed on pseudobulk results using fgsea,^79^ ranking genes by log_2_ fold-change × −log_10_(nominal *P* value), and testing against MSigDB Hallmark, KEGG, Reactome, and GO:BP gene sets and WikiPathways. Normalized enrichment scores and BH-adjusted *P* values were retained for each pathway, cell type, and database.

For astrocyte co-expression network analysis, astrocyte gene counts were aggregated by mouse (*n* = 16), TMM-normalized, and converted to logCPM. Low-expression, ribosomal, mitochondrial, and cross-cell-type RNA carryover genes were excluded. Genes were required to have mean astrocyte expression >5 CPM, and the 2,500 most variable remaining genes were selected. A signed Spearman correlation network was constructed by retaining each gene’s 15 strongest positive correlations > 0.20. Edge stability was assessed using 100 bootstrap iterations, and stable modules were identified by Leiden clustering. Module scores were calculated from the original normalized expression data and tested using a linear model including treatment, sex, and parental origin of the 5xFAD transgene as covariates, with Benjamini-Hochberg correction across modules. Modules were annotated using pathway enrichment and hub genes.

### Mouse behavioral experiments

7–8-month-old male and female *APOE4*;5xFAD mice were randomly assigned to receive rLPs or saline control treatment, with littermates distributed across groups where possible. Behavioral testing began 7 days after intracisternal injection as described above, and the novel object recognition test was performed 10 days after injection. On day 7, mice were habituated to the behavioral testing room for 1 h. On day 8, mice underwent a 10-min open-field test session, which also served as habituation to the testing arena (40 cm L x 40 cm W x 35 cm H). On day 9, mice were exposed to two identical wooden blocks (Premium wooden building blocks set, Cubbie Lee) for 10 min during the familiarization phase. On day 10, one of the familiar objects was replaced with a novel object for a 10-min test session. All behavioral sessions were video recorded and EthoVision software was used for tracking. Treatment groups were blinded when object exploration time was scored. Exploration was defined as the mouse actively investigating an object, as indicated by its nose being oriented toward and directed at the object; time spent near the object without active investigation was not counted. The discrimination index (DI) was calculated as: DI = (*t*_novel_ - *t*_familiar_) / (*t*_novel_ + *t*_familiar_), where *t*_novel_ and *t*_familiar_ represent the time spent exploring the novel and familiar objects, respectively. A DI of zero indicates equal exploration of the two objects, whereas a positive DI indicates preferential exploration of the novel object. Total object exploration time was calculated as the sum of the time spent exploring both objects during the test phase.

## AI Disclosure

Proofig AI, Anthropic’s Claude (Sonnet 5.5), and OpenAI’s ChatGPT (GPT-6.1) were used during manuscript preparation to detect grammatical/typographical errors and accidental figure duplication, and iterate for text clarity. Anthropic’s Claude Code (Opus 5.5) was used for coding checks and assistance.

## Data Availability

Human lipidomics datasets generated in this study will be deposited in the AD Knowledge Portal (Synapse) and made available upon publication, subject to applicable data access requirements. Mouse lipidomics and RNA-seq datasets will be deposited in appropriate repositories. Additional data supporting the findings of this study are available from the corresponding authors upon reasonable request.

## Code Availability

Custom code used for data analysis will be made publicly available on GitHub upon publication.

## Acknowledgments

We thank the ROSMAP study individuals who donated blood and postmortem brain and cerebrospinal fluid samples, and their families. ROSMAP is supported by P30AG10161, P30AG72975, R01AG15819, R01AG17917, U01AG072572, and U01AG046152. ROSMAP resources can be requested at https://www.radc.rush.edu. Thank you to H. Xu in the lab of M. Lehtinen for demonstrating the mouse CSF extraction procedure; E. Rousseau in the lab of M. Cima for exploring the adaptation of the NanoPump setup for CSF extraction; M. Zoller and A. Seguin for exploring additional *in vivo* characterization experiments; T. Ko for stem cell support; U. Geigenmüller and J. M. Bonner for helpful project discussions; U. Geigenmüller, G. Scarpa, and E. Smith for feedback on the manuscript; Y. Zhou, T. Garvey, and M. Mazzanti for administrative support; K. Lee for mouse colony support. R. Pinals acknowledges support from the Schmidt Science Fellows program in partnership with the Rhodes Trust and the Burroughs Wellcome Fund Career Award at the Scientific Interface (CASI). Y. Koh was supported by NIH/NIA Award K00AG083111. We thank the generous support of the WoodNext Foundation, the Belfer Neurodegeneration Consortium, the Robert and Renee Belfer Family Foundation, the Freedom Together Foundation, the Cure Alzheimer’s Fund, the BT Charitable Foundation, Chejin Lee, and the McNamara Manthripragada Fund. This work was supported in part by the Koch Institute Support (core) Grant P30-CA14051 from the National Cancer Institute. We thank the Koch Institute’s Robert A. Swanson (1969) Biotechnology Center for technical support specifically Peterson (1957) Nanotechnology Materials Core Facility (RRID:SCR_018674). Graphic illustrations were generated using BioRender.

## Author Contributions

R.L.P. and L-H.T. conceived and supervised the project; R.L.P., Y.K., A.T., C.F.L.C., D.v.M., and L.A. designed and performed the experiments and analyses, with assistance from I.G., A.B., C-Y.Y., A.N., L.B.C., J.W., P.B., D.L., R.H., A.C., I.C., E.K., D.R.-A., E.M., K.G., J.C., and F.L.; Z.K. performed sequencing data analyses and visualization; A.D. and K.P. performed lipidomics on mouse samples and C.C. supervised; D.S.Y. performed electron microscopy; D.A.B. provided MAP cohort samples; R.L.P., Y.K., and L-H.T., wrote the manuscript with input from authors.

---

**Extended Data Figure 1.**
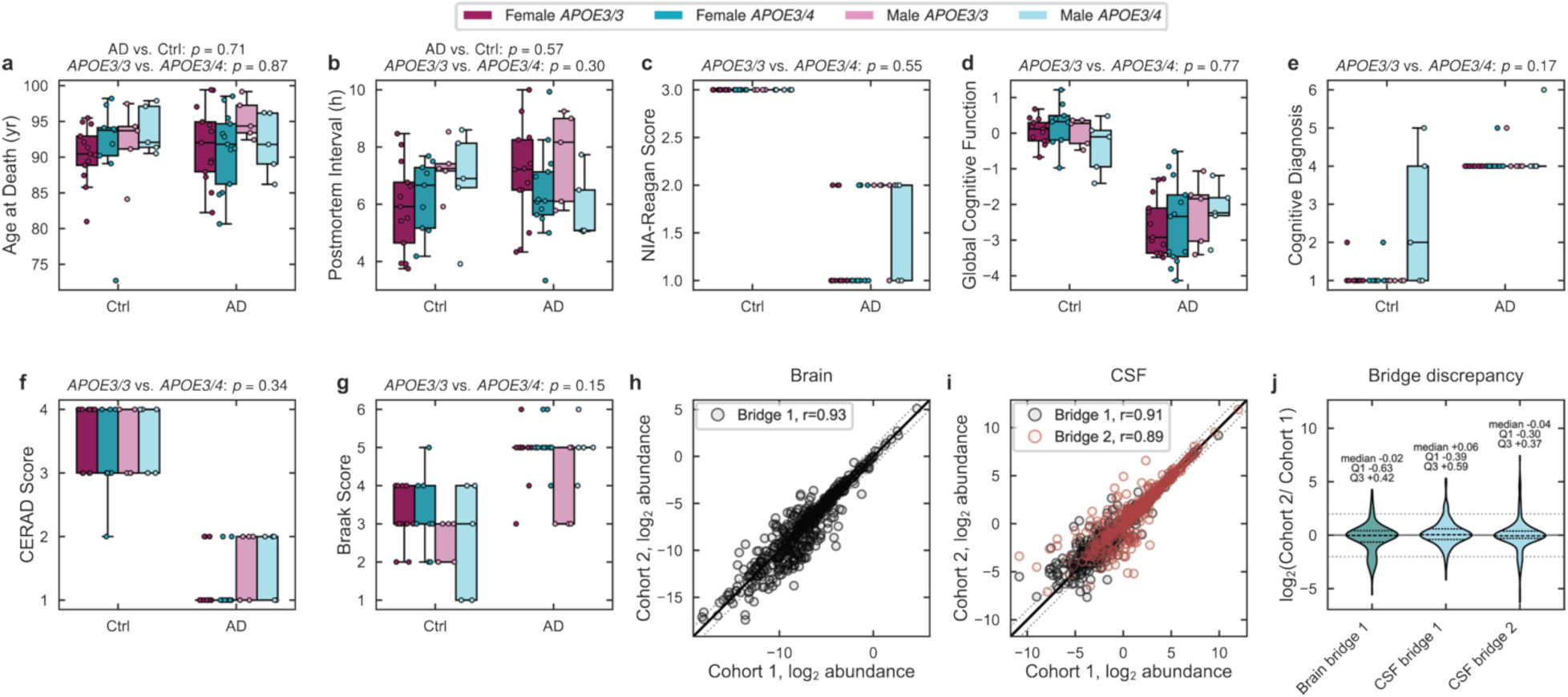
Human MAP cohort design across AD status, *APOE* genotype, and chromosomal sex. Cohort metadata for individuals without (Ctrl) or with neuropathologic Alzheimer’s disease (AD) diagnosis by NIA-Reagan criteria, *APOE3/3* (magenta) and *APOE3/4* (cyan) genotypes, and females (dark) and males (light) showing **(a)** age at death, **(b)** postmortem interval, **(c)** NIA-Reagan diagnosis, **(d)** global cognitive function at last visit, **(e)** final consensus cognitive diagnosis, **(f)** neuritic plaques (CERAD score), and **(g)** neurofibrillary pathology (Braak stage). Pairwise comparisons of diagnoses (AD vs. Ctrl) and genotype (*APOE3/3* vs. *APOE3/4*) were not significantly different for age and postmortem interval **(a,b)**, nor were pairwise comparisons of genotype for different neuropathologic and cognitive measures **(c–g)**. Statistical significance was determined using a two-sided Mann-Whitney U test. Bridge samples analyzed across both mass spectrometry batches showed strong agreement in log_2_ lipid abundances for **(h)** one PFC sample and **(i)** two CSF samples (Pearson’s *r* shown). Note the second PFC sample was removed during quality-control filtering due to injection volume error. **(j)** Distribution of cross-batch discrepancy for bridge samples, calculated as log_2_(Cohort 2/Cohort 1) abundance for each aligned lipid feature; median and interquartile ranges are indicated.

**Extended Data Figure 2.**
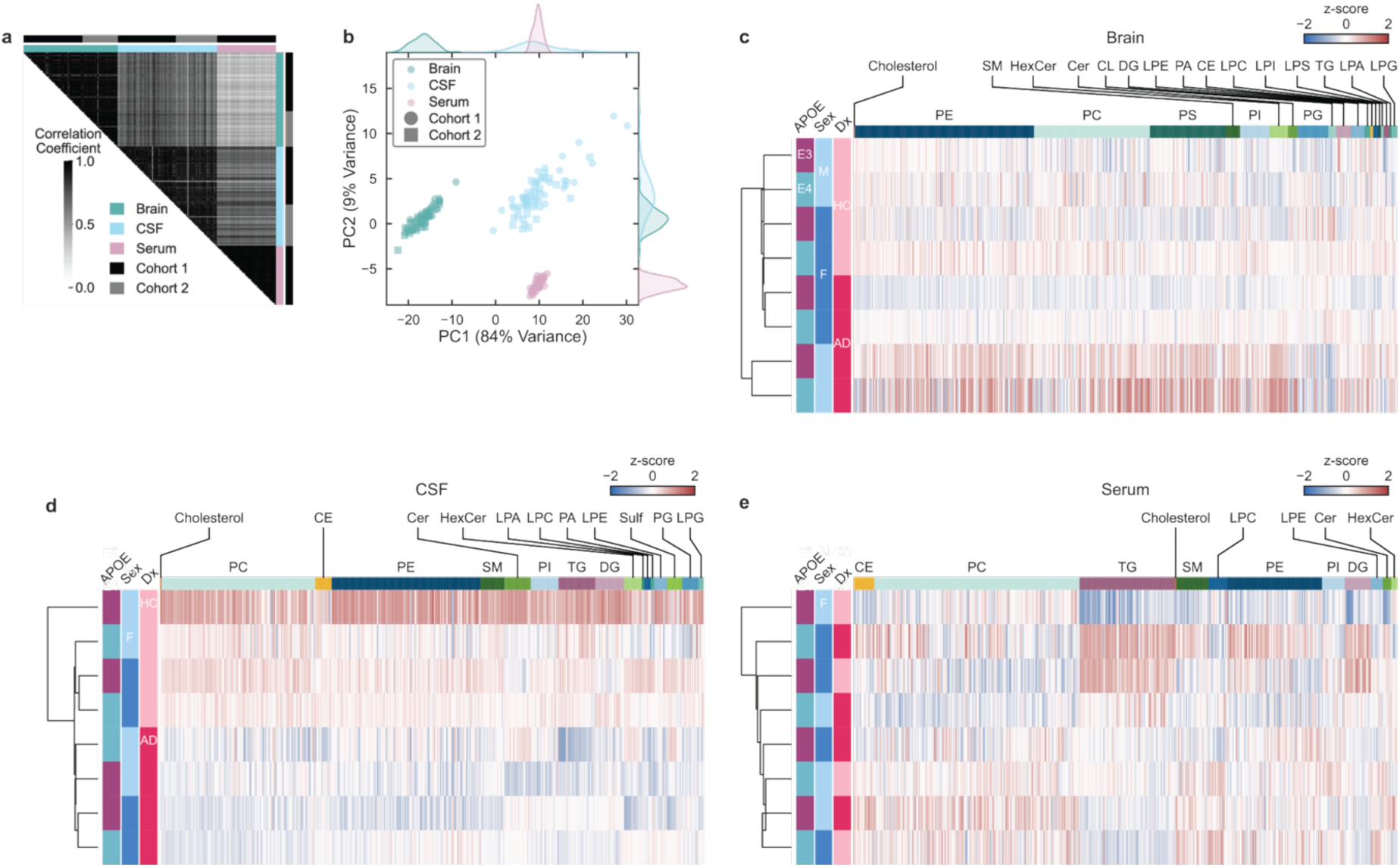
Human MAP cohort lipidomics overview. **(a)** Heatmap of Pearson correlation coefficients between samples using log_2_-transformed lipid abundances of 215 lipid species aligned across brain (green), CSF (blue), and serum (pink). **(b)** Principal component analysis (PCA) of normalized (z-score of log_2_-transformed) lipid abundances in brain, CSF, and serum, restricted to the 215 aligned lipid species with complete measurements across all samples. Heatmap of z-scores of **(c)** brain, **(d)** CSF, and **(e)** serum log_2_-transformed lipid abundance (columns) ordered by lipid class fractional abundance in each tissue or biofluid, averaged across individuals in each sub-population (rows), grouped using a dendrogram generated by hierarchical clustering based on normalized lipid abundance profiles with Euclidean distance and median linkage. Block colors correspond to negative (dark blue), zero (white), and positive (dark red) z-score saturated at ±2. Ether-linked lipids are grouped with the corresponding headgroup lipid class.

**Extended Data Figure 3.**
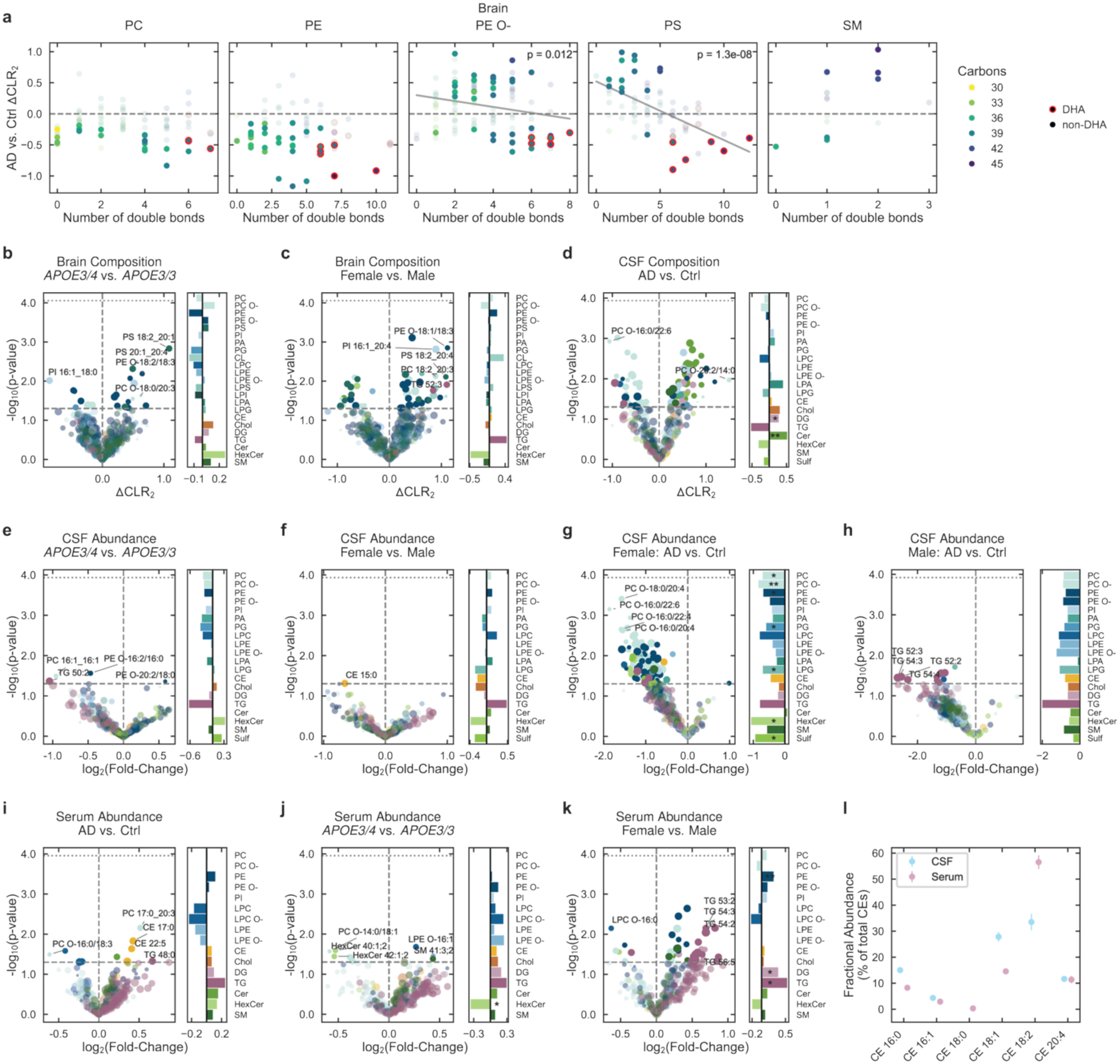
Human MAP cohort brain, CSF, and serum lipidomics. **(a)** Difference in CLR_2_ lipid abundance for AD vs. Ctrl brain as a function of total number of double bonds, for different lipid classes per panel. Markers indicate individual lipid species and are colored by total acyl-chain carbon number; DHA-containing lipid markers are outlined in red. Significantly changing lipids are shown with darker opacity. Gray lines denote the linear fit across all lipids, with annotated *P* value where significant. Volcano plots (-log_10_ *P* value vs. log_2_ fold-change) of lipids detected in **(b–c)** brain, **(d–h)** CSF, and **(i–k)** serum, with comparisons listed above each sub-panel. Markers in volcano plots indicate lipid species, grouped by color for different lipid classes as defined on the right-side bar plots. *P* values were determined using a two-sided *t*-test assuming equal variances. Short-dashed line indicates FDR-adjusted *P* = 0.05 (Benjamini-Hochberg correction). **(l)** Fractional abundance of CE species in CSF (blue) and serum (pink) represented as the percent of total CEs. Error bars denote 95% confidence intervals of the mean using a t-statistic.

**Extended Data Figure 4.**
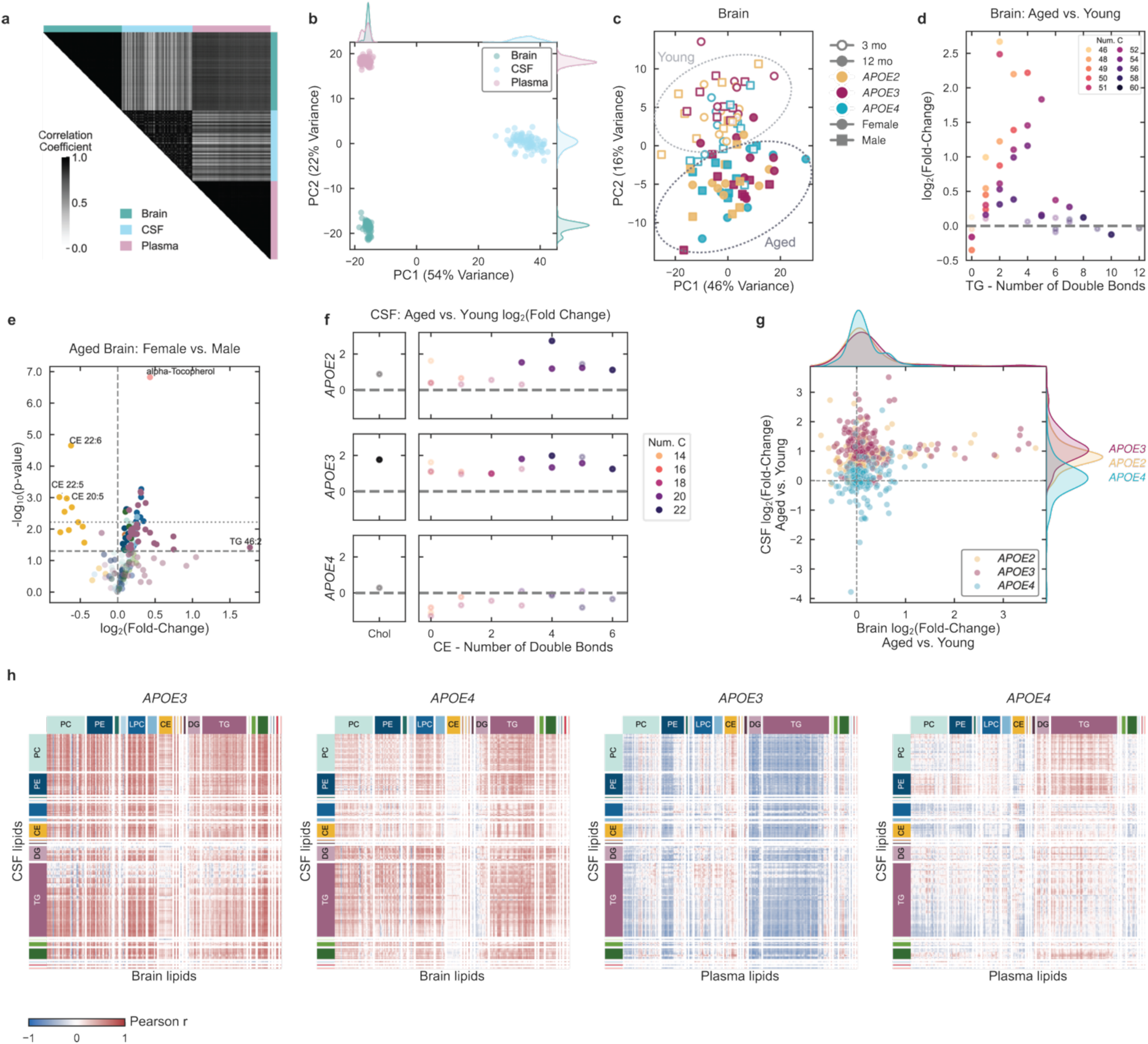
Mouse cohort design and lipidomics overview. **(a)** Heatmap of Pearson correlation coefficients between samples using log_2_-transformed abundances of 1071 lipid features aligned across brain (green), CSF (blue), and plasma (pink). **(b)** Principal component analysis (PCA) of aligned, normalized (z-score of log_2_-transformed) lipid abundances in brain, CSF, and plasma. **(c)** PCA of normalized brain lipid abundances for each age (3 months: open, 12 months: filled), *APOE* genotype (*APOE2*: yellow, *APOE3*: magenta, *APOE4*: cyan), and sex (female: circles, male: squares). Brain lipidomic fingerprints primarily cluster by age, with dashed outlines drawn to guide the eye. **(d)** Log_2_ fold-change in TG abundance for aged vs. young brain as a function of the total number of double bonds. Markers indicate individual TG species and are colored by the total number of carbons in the acyl chains. Significantly changing species (two-sided *t*-test, *P* < 0.05) are shown with darker opacity. **(e)** Volcano plot of lipids in the aged brain for female vs. male mice, revealing decreased CEs and increased TGs. Short-dashed line indicates FDR-adjusted *P* = 0.05 (Benjamini-Hochberg correction). **(f)** For each *APOE* genotype, log_2_ fold-change in CSF cholesterol (left) and CE (right) abundance in aged vs. young mice. CE species are arranged by total number of double bonds and colored by the total number of carbons in the acyl chains. Significantly changing species (two-sided *t*-test, *P* < 0.05) are shown with darker opacity. **(g)** Log_2_ fold-changes in lipid abundances for aged vs. young mice in CSF vs. brain show increased brain lipid abundance with age across *APOE* genotypes, accompanied by increased CSF lipid abundance in *APOE2* and *APOE3*, but not *APOE4*, mice. Corresponding PDFs by KDE are included on the top and side panel for brain and CSF, respectively. **(h)** Heatmaps of Pearson correlation coefficients between z-scores of CSF lipids as rows and brain lipids (first two panels) or plasma lipids (second two panels) as columns, grouped by color for different lipid classes in *APOE3/3* (first and third panels) and *APOE4/4* (second and fourth panels), colored from red (r = 1) to white (r = 0) to blue (r = -1).

**Extended Data Figure 5.**
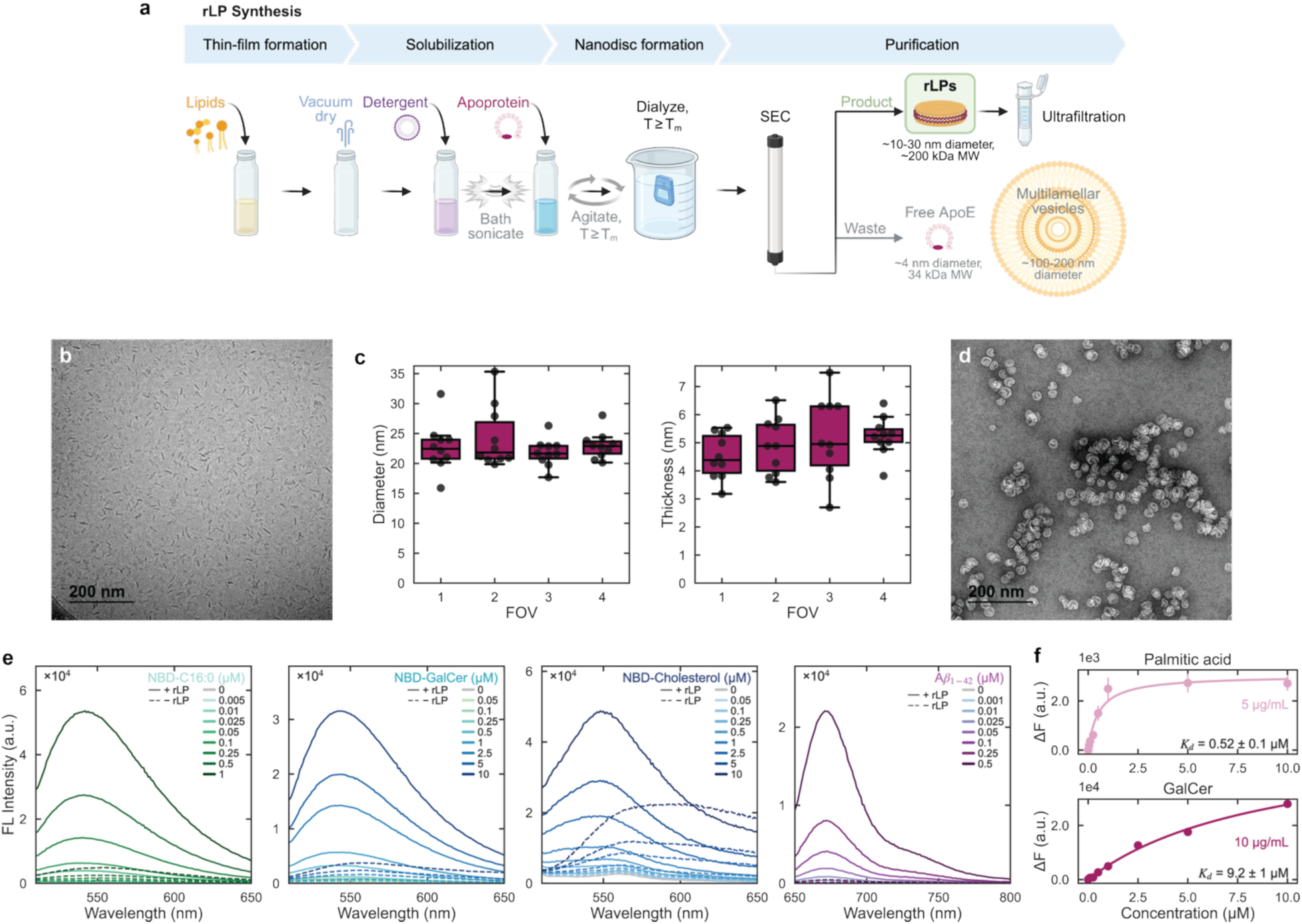
rLP synthesis and characterization. **(a)** Schematic of rLP synthesis by thin-film hydration followed by ApoE incorporation and purification by size-exclusion chromatography (SEC) and ultrafiltration. **(b)** Representative cryogenic transmission electron microscopy (cryoTEM) image of discoidal rLPs in a frozen-hydrated state. **(c)** Quantification of rLP diameter (long axis, left) and bilayer thickness (right) from TEM images of one rLP preparation (10 particles measured in each of 4 fields of view). Boxes show median and interquartile range; whiskers extend to 1.5× the interquartile range; points are individual particles. **(d)** Representative TEM image of lipid particles formed in the absence of ApoE. **(e)** Fluorescence emission spectra of nitrobenzoxadiazole (NBD)-labelled palmitic acid (NBD-C16:0), galactosylceramide (GalCer), and cholesterol, or HiLyte Fluor 647-tagged amyloid-β peptide (Aβ42), incubated at varying concentrations with (solid lines) or without (dashed lines) rLPs. Spectra represent the mean of *n* = 3 wells per condition. **(f)** Fluorescence responses for NBD-C16:0 (top) and NBD-GalCer (bottom) with 5 or 10 µg/mL rLPs, respectively. Data are presented as mean ± SEM (*n* = 3); solid lines indicate fitted binding isotherms, with apparent dissociation constants (K_D_) and fitting standard errors indicated.

**Extended Data Figure 6.**
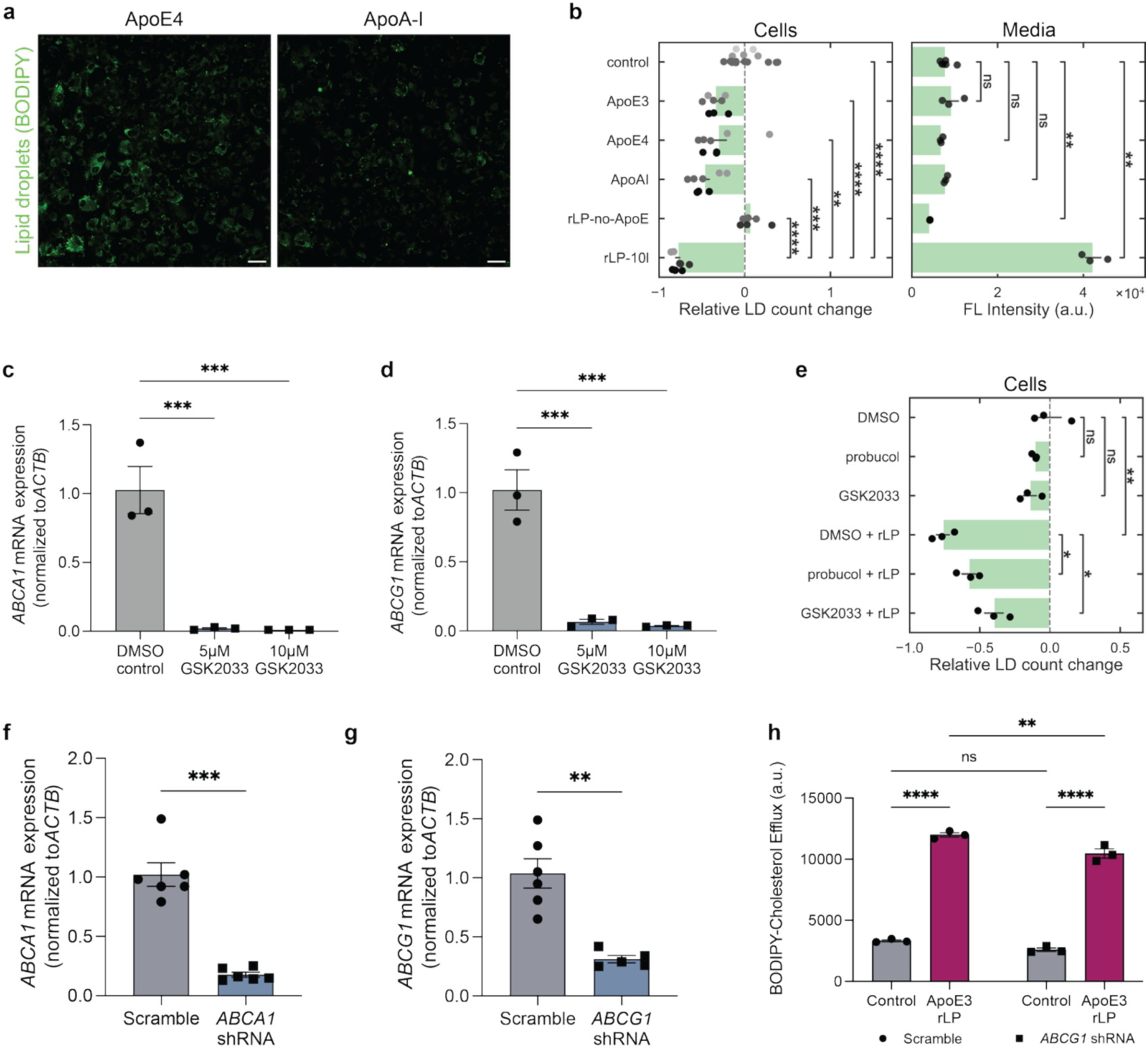
rLP functional testing with cellular efflux assays. **(a)** Representative images of BODIPY-cholesterol-loaded *APOE4* astrocytes treated with free ApoE4 or ApoA-I. **(b)** Relative lipid droplet (LD) count change in cells (left) and BODIPY-cholesterol fluorescence in medium (right) following treatment with no acceptor, free ApoE3, free ApoE4, free ApoA-I, ApoE-free rLPs, or full rLPs (each at 10 µg/mL by protein mass; ApoE-free particles were diluted by the same factor as full rLPs). Markers denote individual wells (LD counts averaged over *n* ≥ 3 imaged regions per well), and marker color (left) denotes independent experimental batches. **(c–d)** RT-qPCR validation of GSK2033 reduction of **(c)** *ABCA1* and **(d)** *ABCG1*, normalized to *ACTB*. **(e)** Relative LD count change in cells following pretreatment with DMSO (vehicle), probucol (20 µM), or GSK2033 (10 µM), with or without subsequent rLP treatment, with markers as described in **(b)**. **(f–g)** RT-qPCR validation of shRNA-mediated knockdown of **(f)** *ABCA1* and **(g)** *ABCG1*, normalized to *ACTB* and scrambled-shRNA controls. **(h)** BODIPY-cholesterol efflux into medium from *APOE4* astrocytes expressing scrambled (left) or *ABCG1* (right) shRNA, treated with no acceptor or rLPs. **(b, e)** Error bars denote 95% confidence intervals of the mean using a t-statistic. Statistical significance was determined by Welch’s two-sided t-test. **(c–d, f–h)** Data are presented as mean ± SEM (*n* ≥ 3). Statistical significance was determined by one-way ANOVA with Tukey’s multiple comparisons. * *P* < 0.05, ** *P* < 0.01, *** *P* < 0.001, **** *P* < 0.0001.

**Extended Data Figure 7.**
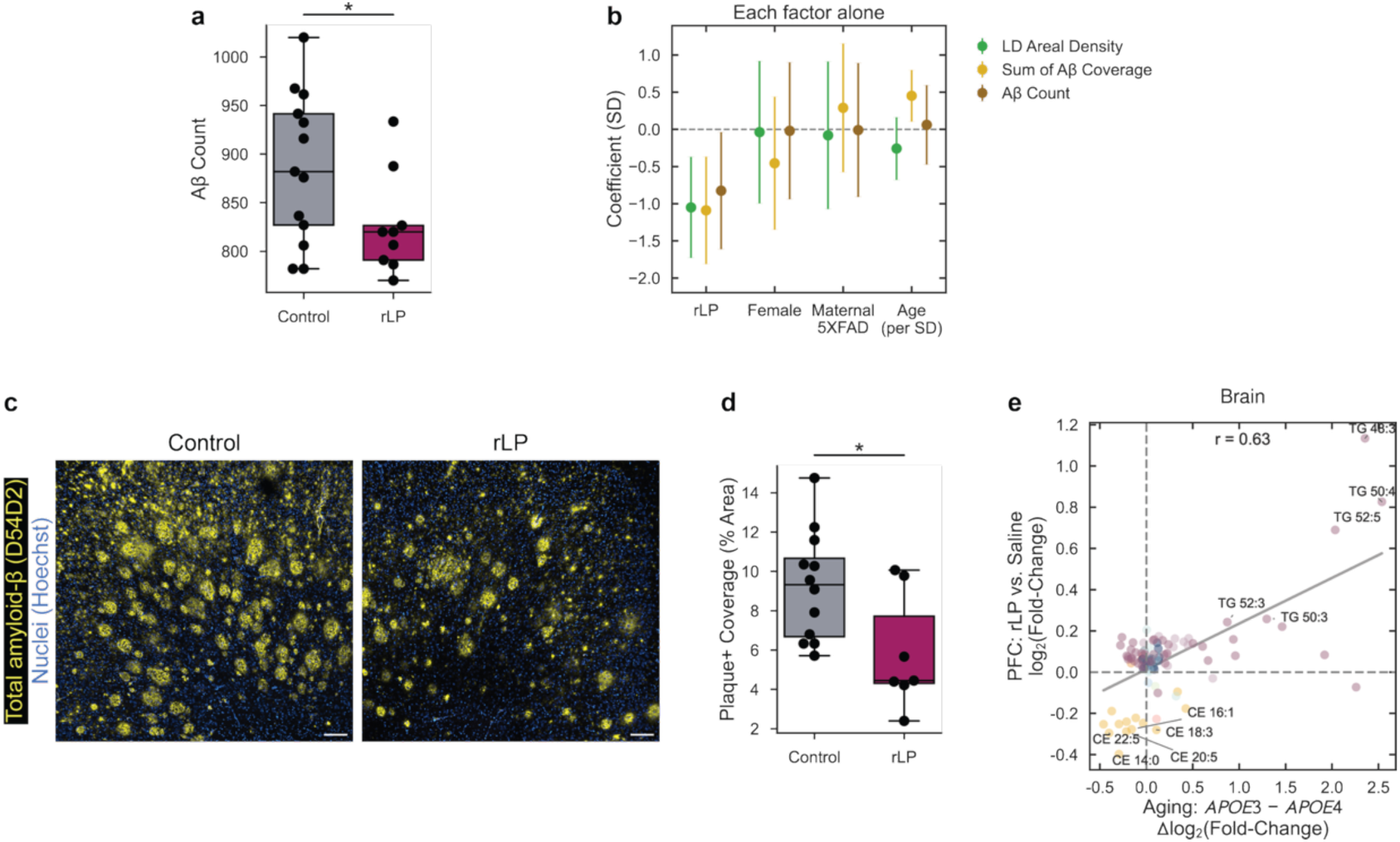
rLP-associated changes in *APOE4;*5xFAD mouse brain pathology and lipids. **(a)** Quantification of Aβ count for saline control vs. rLP treatment. Markers denote individual mice (*n* ≥ 7), averaged over two brain sections per mouse. Statistical significance was determined using Welch’s two-sided t-test. * *P* < 0.05, ** *P* < 0.01, *** *P* < 0.001, **** *P* < 0.0001. **(b)** Standardized univariant regression coefficient of lipid droplet (LD) and Aβ burden z-scores regressed separately against mouse sex, parental inheritance of the 5xFAD transgene, and the z-score of exact age at injection. **(c)** Representative images of D54D2-stained total Aβ (yellow) and Hoechst-stained cell nuclei (blue) in the mouse prefrontal cortex for control (left) and rLP treatment (right). **(d)** Prefrontal cortex area positive for Aβ plaques for control (gray) vs. rLP treatment (magenta). Markers denote individual mice. Statistical significance was determined using Welch’s two-sided t-test. **(e)** Correlation of lipid abundance changes following rLP treatment in the prefrontal cortex (log_2_ fold-change, rLP vs. saline) with *APOE*-dependent aging effects in mouse brain, calculated as the difference between aging-associated log_2_ fold-changes in *APOE3/3* and *APOE4/4* mice. Markers denote individual lipid species, with selected TG (purple) and CE (yellow) species labelled. Pearson correlation coefficient, *r* = 0.63. * *P* < 0.05, ** *P* < 0.01, *** *P* < 0.001, **** *P* < 0.0001.

**Extended Data Figure 8.**
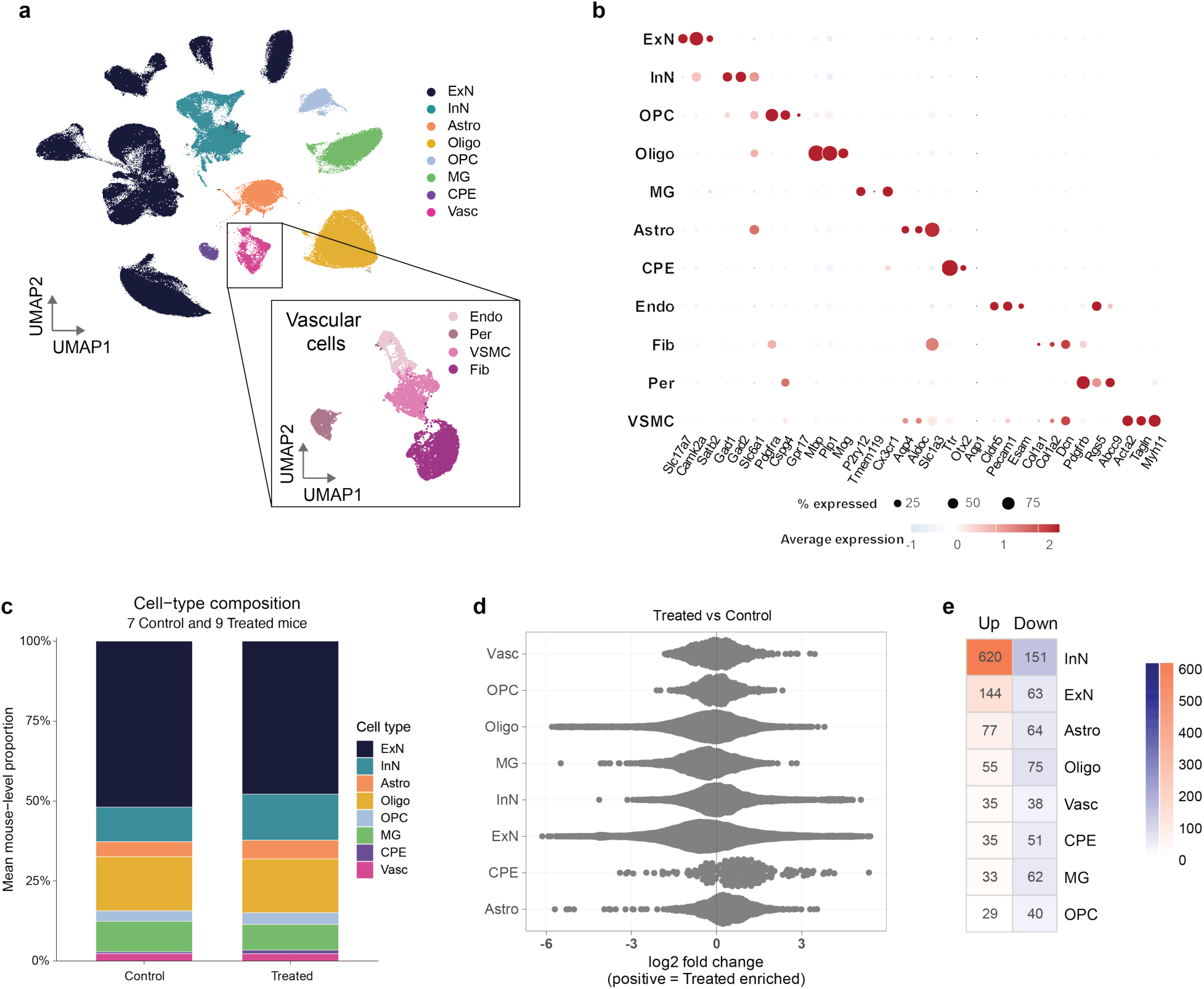
Cell-type composition in rLP-treated mice. **(a)** Cell-type clustering by uniform manifold approximation and projection (UMAP) of 300,399 nuclei after data filtering as outlined in the Methods. **(b)** Dot plot showing expression of canonical marker genes across major neural, glial, choroid plexus and vascular cell types. Dot size represents the percentage of nuclei expressing each gene, and color indicates scaled average expression. **(c)** Mean mouse-level proportions of major cell types in control (*n* = 7) and rLP-treated (*n* = 9) mice. **(d)** Milo differential-abundance analysis comparing rLP-treated and control mice. Each point represents a cellular neighborhood (no neighborhoods reached significance at spatial FDR < 0.1); positive log_2_ fold-changes indicate enrichment following rLP treatment. **(e)** Summary heatmap of number of genes upregulated (red) and downregulated after rLP treatment, by cell type.

**Extended Data Figure 9.**
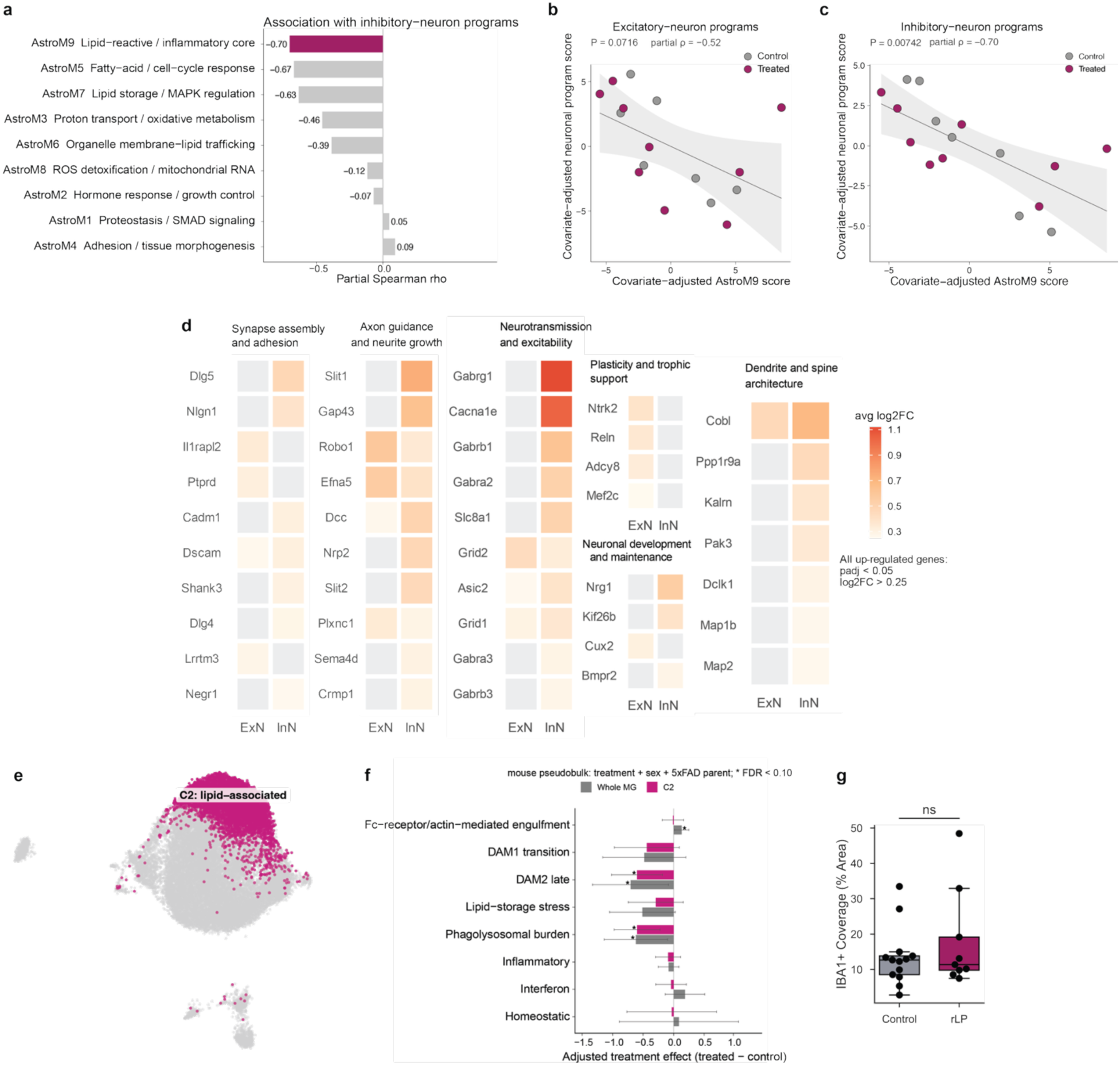
Coordinated glial and neuronal responses in rLP-treated mice. **(a)** Partial Spearman correlation coefficients between astrocyte co-expression module scores and inhibitory-neuron program scores after adjustment for treatment, sex, and parental 5xFAD origin. **(b–c)** Associations of the AstroM9 lipid-reactive/inflammatory module with **(b)** excitatory-and **(c)** inhibitory-neuron programs across mice. Axes show covariate-adjusted module scores; points represent individual mice and are colored by treatment group. Lines and shaded regions indicate linear fits and 95% confidence intervals. Reported values are partial Spearman correlations and nominal P values. **(d)** Selected genes upregulated by rLP treatment in excitatory and inhibitory neurons, grouped by neuronal function. Tile color represents the average log_2_ fold-change; displayed genes met adjusted *P* < 0.05 and log_2_ fold-change > 0.25 in at least one neuronal population. **(e)** Microglial UMAP highlighting the C2 lipid-associated state. **(f)** Adjusted treatment effects on microglial functional programs across all microglia (Whole MG, gray) and within the C2 population (magenta). Models were fitted to mouse-level pseudobulk scores and included treatment, sex, and parental 5xFAD origin; negative coefficients indicate lower program activity following treatment. Error bars indicate 95% confidence intervals. **(g)** Prefrontal cortex area positive for IBA1 for saline control (gray) vs. rLP treatment (magenta). Markers denote individual mice. Statistical significance was determined using Welch’s two-sided t-test. Abbreviations: ExN, excitatory neurons; InN, inhibitory neurons; OPC, oligodendrocyte precursor cells; Oligo, oligodendrocytes; MG, microglia; CPE, choroid plexus epithelium; Endo, endothelial cells; Fib, fibroblasts; Per, pericytes; VSMC, vascular smooth-muscle cells.

**Extended Data Figure 10.**
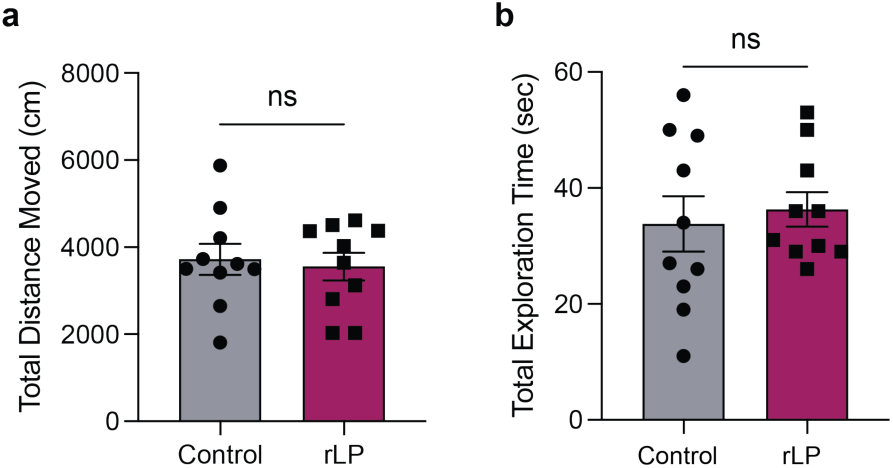
Similar locomotor activity during novel object recognition test. Quantification of **(a)** total distance moved during the test session (10 minutes) and **(b)** total exploration time did not differ between saline control (gray) and rLP treatment (magenta) group. Unpaired t-test for both quantification; ns, not significant.

